# Transmission mode shapes host specialization of the phyllosphere microbiome

**DOI:** 10.1101/2023.05.08.539874

**Authors:** Kyle M. Meyer, Isabella E. Muscettola, Ana Luisa S. Vasconcelos, Julia K. Sherman, C. Jessica E. Metcalf, Steven E. Lindow, Britt Koskella

**Affiliations:** Department of Integrative Biology, University of California, Berkeley, Berkeley, USA 94720; Department of Soil Science, College of Agriculture “Luiz de Queiroz”, Universidade de São Paulo, Piracicaba, Brazil 13418-900; Department of Ecology and Evolutionary Biology, Princeton University, Princeton, NJ, USA 08544; Department of Plant and Microbial Biology, University of California, Berkeley, Berkeley, CA, USA 94720; Chan Zuckerberg Biohub, San Francisco, CA 94158, USA

**Keywords:** Microbiome transmission, Microbiome specialization, Experimental evolution, Phyllosphere, Microbiome engineering

## Abstract

The collection of microorganisms inhabiting aboveground plant tissue, termed the phyllosphere microbiome, is shaped by both microbial dispersal and host filtering effects. Because plants often occur in multispecies assemblages, microbiome diversity is likely shaped by plant community composition through microbial transmission between conspecific (same species) and heterospecific (different species) host plants. Although rarely examined, the relative incidence of con-versus heterospecific transmission should impact microbiome-level specialization. Using four species of plants we experimentally tested this idea by passaging an initially diverse microbial community either between conspecific or heterospecific plants in the greenhouse. While conspecific transmission lines exhibited persistent host effects, these effects decreased in the heterospecific transmission lines, suggesting a shift towards more generalist microbiomes. Similarly, when microbiomes were transplanted onto a set of novel host plant species, host effects were weaker for these heterospecific lines than the conspecific lines. Finally, microbiomes conspecifically passaged on tomato plants were found to outcompete those passaged on either bean or pepper when co-inoculated onto tomato hosts, suggesting microbiome-level host specialization under conspecific transmission. Overall, we find that both transmission mode and host association history shape microbiome diversity, that repeated conspecific transmission results in microbiome specialization, and that repeated heterospecific transmission can drive microbiomes to develop generalist characteristics.

## Introduction

Transmission mode, i.e. the means by which microorganisms move between hosts, has the potential to shape host-microbe interactions over both short, microevolutionary, and longer, macroevolutionary, timescales (1, 2). Both theory and empirical evidence suggest that strict vertical (parent-to-offspring) transmission of individual microbial symbionts or pathogens has the potential to drive highly specialized adaptations for the symbiont, including genome reduction and streamlined metabolic capability (3), as well as reduced virulence (4, 5). Similarly, transmission between conspecific (same species) hosts has the potential to drive the formation of mutualistic relationships, whereby the fitness of both participants increases through the shared benefits of cooperation (6). On the other hand, horizontal, or heterospecific, transmission (i.e. between unrelated hosts) can drive increased virulence in pathogens (7), and in symbionts may promote a “jack-of-all-trades, master of none” strategy whereby symbionts are less beneficial to host fitness (6), less likely to gain host-specific adaptations and more likely to return to autonomy (8). Importantly, much of this framework has centered on pairwise interactions, and a considerable gap remains regarding whether transmission mode may also shape the local adaptation of the complex consortia of microorganisms residing in or on a host, i.e. the microbiome.

Aboveground plant tissue, also known as the phyllosphere, harbors a distinct microbiome comprised of bacteria, fungi, and viruses (9, 10). Phyllosphere microbiomes are implicated in multiple facets of plant health including disease resistance (11–13), auxin production (14), and primary productivity (15). These microbial communities assemble primarily *de novo* at seed germination or leaf emergence and tend to follow distinct assembly patterns depending on the host plant species (16). These host species identity effects arise on leaf surfaces through ecological (host) filtering resulting from different chemical and physical features of the host (17, 18), as well as immune activity (19, 20), molecular signaling (9), and tissue/barrier formation (21). Importantly, such filtering effects depend on the diversity of microbial taxa arriving on the leaf, and are thus reliant on the process of dispersal (22).

A major source of microbial dispersal into the phyllosphere is the nearby vegetation (23); and the species identity, size, and environmental context of this vegetation has been shown to impact microbiome assembly (24–26). In low diversity plant communities, we might expect transmission among related plants to be more likely than among unrelated plants, and this consistent plant-driven selective pressure (ecological/plant filtering effect) could promote host specialization of microbiomes. In contrast, frequent transmission among different plant species, as might be the norm in higher diversity plant communities, may disrupt microbiome specialization and instead select for taxa that are able to persist through multiple sets of host filters, driving increased microbiome generalism. Better understanding the means by which specialization or generalism arise could inform new strategies in sustainable agriculture, conservation, and microbiome engineering (27–32). The longer-term consequences of such differences in transmission could also shed light on the formation or maintenance of symbiotic relationships over evolutionary time.

To better understand how microbiome transmission might shape host specialization, we experimentally passaged a diverse microbial community initially derived from field-grown tomato plants on either conspecific (one of four plant species consistently throughout passaging) or heterospecific (alternating between tomato and each of the other three species at each passage) plant hosts. We used 16S rRNA gene amplicon sequencing to track microbiome assembly across treatments over six plant passages, and then further explored specialization of the microbiomes in two follow-up common garden experiments, one that applied experimental community coalescence to directly compete microbiomes, and another in which microbiomes were transplanted onto novel hosts. In both the primary passaging experiment and follow up studies, we find clear but nuanced evidence of microbiome specialization, as well as clear plant filtering effects upon this common pool of microbial taxa. These findings make clear that by altering the species pool on which host selection acts, transmission mode can impact the outcome of microbiome assembly and shape the prospects for host specialization of the microbiome.

## Methods

### Experimental design - Passaging experiment

To generate the initial community for selection, we created a diverse microbial inoculum by sampling epiphytic (leaf surface) microbiomes from tomato leaves collected in two regions of California at separate times. The first collection was at Coastal Roots Farm, located in Encinitas, CA in May 2019, where plants were in early vegetative growth (pre-flowering) and had been established in the field for roughly 4 weeks. The second collection was at the University of California - Davis student organic farm in August 2019, where plants were 3-4 months old and in the fruit ripening stage. In both cases, leaves were excised from the plants, placed into sealable sterile 1-gallon plastic bags (Ziplock, USA), and transported in a chilled cooler to the laboratory and held at 4° C. The day following collection, epiphytic microbial members of the phyllosphere were gently dissociated from leaf surfaces using a sonicating water bath (Branson 5800) using a ratio of roughly 0.6 g of leaf material to 1 ml 10 mM MgCl_2_. In total, 4.7 kg of leaf material were processed from the Encinitas collection and 2.4 kg from the Davis collection. Leaf wash was pelleted by centrifuging at 4000 rcf and 10° C for 10 minutes and then resuspended in KB broth and mixed at a 1:1 ratio with glycerol for preservation at −20° C. The leaf wash from both collections was combined to serve as the initial inoculum for the experiment. Prior to combining, aliquots of each collection were set aside to later test whether they differed in establishment ability.

We used four plant hosts for passaging: cultivated tomato (*Solanum lycopersicum var.* Moneymaker), *Solanum pimpinellifolium* (tomato wild ancestor), bell pepper (*Capsicum anuum var.* Early Cal Wonder), and bush bean (*Phaseolus vulgaris* var. Bush Blue Lake 274). Due to differences in germination and development rate, seed planting times were adjusted so that plants were roughly equivalent in size at initial inoculation. Tomato and *S. pimpinellifolium* were 4.5 weeks post seed planting, peppers were 5.5 weeks, and beans were 2.5 weeks.

To begin the passaging experiment, 6 plants from each of the four hosts was inoculated with the same initial inoculum (Fig. 1A,B), which was prepared by thawing the freezer stock, centrifuging at 4000 rcf and 10° C for 10 minutes to pellet cells, decanting the supernatant, and re-suspending cells in sterile 10 mM MgCl_2_. For the first week of inoculation, inoculum volumes per plant were 4 ml for tomato, *S. pimpinellofolium*, and bean plants, and 3 ml for the slightly smaller pepper plants. Inocula were sprayed onto the adaxial (top) and abaxial (bottom) sides of leaves using ethanol- and UV-sterilized hand-held misters. After inoculation, the moist, sprayed plants were placed in a chamber maintaining ca. 100% relative humidity for 20 hours in order to maintain leaf moistness, thus encouraging microbial growth, before being transferred to a greenhouse. In both cases, plant location on the bench was randomized among treatments. Because of both space limitations in the misting chamber, and the scale of inoculations, we divided the plants into 2 blocks, such that block 1 contained replicates 1-3 and the blank (no inoculum) controls, all of which were sprayed on one day, while block 2 contained replicates 4-6, all of which were sprayed 2 days later. Plants from experimental blocks 1 and 2 were placed on different adjacent benches. A week later plants received a second inoculation following the same procedure, but with slightly higher inoculum volumes: 5 ml per plant for tomato, *S. pimpinellifolium*, and bean plants and 3.5 ml for pepper plants. The plants were then returned to the greenhouse for another week. 1 week after the final inoculation (2 weeks since the first inoculation), leaf material from each plant was excised using ethanol-sterilized scissors, placed in separate sterile plastic bags (Ziplock, USA), and transported in a chilled cooler to the laboratory where it was then weighed and combined with 200 mL sterile 10 mM MgCl_2_ and sonicated as described above. Cells from each microbiome sample were then pelleted and preserved using the above-described centrifugation procedure.

**Fig. 1:**
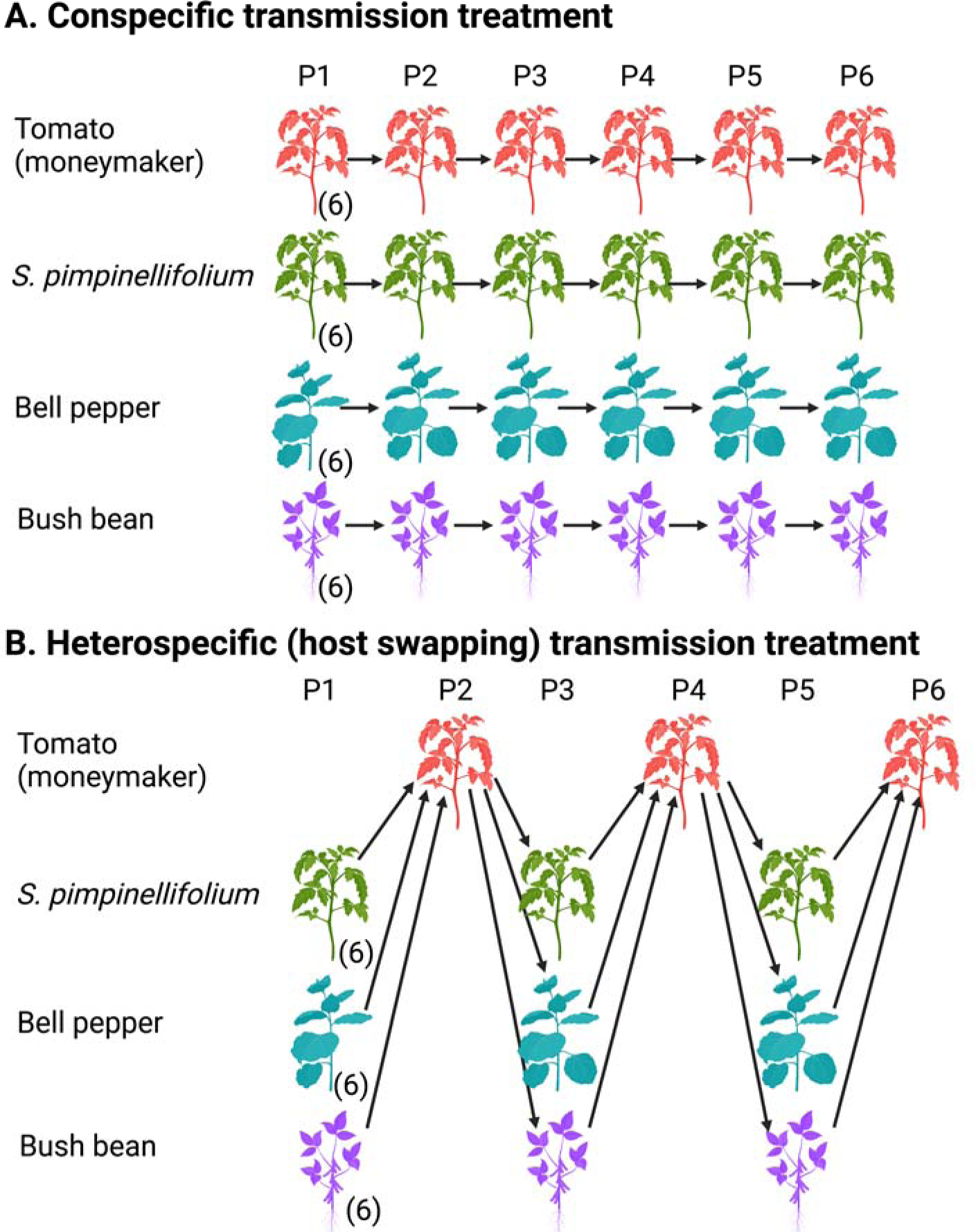
Experimental design of the main passaging experiment where a diverse microbial inoculum was sprayed on to 4 species of plant (moneymaker tomato, *Solanum pimpinellifolium*, bell pepper, and bush bean), allowed to establish and grow, and then be harvested to act as inoculum for the subsequent passage (P2-6). A) Conspecific (same species) transmission treatment whereby replicate microbiome lineages are passaged only on plants of the same species. B) Heterospecific, or host swapping, transmission treatment whereby replicate lineages start on *S. pimpinellifolium*, bell pepper, or bush bean host plants but passage back and forth between tomato hosts and host species of origin. In both cases, each replicate line remained independent across passages (i.e. microbiomes from one individual plant were used as inoculum for one plant in the next passage).

During each subsequent passage, experimental plants were inoculated once per week for 3 weeks, followed by a week in the greenhouse. Independent microbiome lines were maintained for every experimental replicate. Inocula were prepared at the end of each passage in the same way as described above for the initial inoculation. The inoculum volume applied increased successively from week 1 to 3 to account for plant growth. The experiment involved 2 treatment types: 1) conspecific transmission, whereby 6 replicate community lines were passaged between members of the same plant species for 6 successive passages (Fig. 1A), and 2) heterospecific transmission (i.e. host swapping), whereby 6 replicate lines of communities started on non-tomato plants (i.e *S. pimpinellifolium*, pepper, and bean) were inoculated onto tomato plants every other passage (i.e. for passages 2, 4, and 6), thus disrupting conspecific transmission (Fig. 1B). The two experimental controls were: 1) blank inoculum, wherein sterile 10 mM MgCl_2_ was sprayed onto 1 plant of each species at each passage in order to survey the diversity and identity of microbes that might establish in the greenhouse environment, and 2) a seasonality control, in which an aliquot of the combined initial inoculum was sprayed onto 3 replicate tomato plants in order to ask whether similar initial selection results were obtained at each passage throughout the experiment. Each passage thus involved 49 plants, 24 of which were conspecific transmission lines (6 tomato, 6 *S. pimpinellifolium*, 6 pepper, and 6 bean), 18 of which were heterospecific lines (*S. pimpinellifolium*-tomato, pepper-tomato, and bean-tomato), and 7 were controls (4 blank and 3 abiotic). The six experimental passages took place between September 2019 and July 2020, with every plant of each replicate lineage surviving.

### Community coalescence experiment to examine microbiome specialization after final passage

To test for microbiome host specialization, we devised a community coalescence experiment involving the microbiome lines that were conspecifically passaged on tomato, pepper, or bean hosts. For each plant species, the six replicate lines were combined into a single inoculum. The live bacterial cells in each of these 3 groups were quantified using PMA treatment followed by droplet digital PCR (ddPCR) targeting cells with 16S rRNA gene copies (33, 34). Each week for 3 weeks these groups were combined, quantified, and adjusted to the lowest concentration needed to equalize live bacterial cell numbers. Equalized live cell totals were 1.82 x 10^5^, 1.38 x 10^5^, and 1.44 x 10^5^ cells for weeks 1, 2, and 3, respectively. Coalescence treatments were devised by combining the tomato and pepper groups and the tomato and bean groups at a 1:1 ratio of live cells. Following equalization, each group was split into 6 tubes and adjusted to 3.5 ml total volume using sterile 10 mM MgCl_2_ to serve as replicate inocula at roughly 8.6 x 10^3^, 6.6 x 10^3^, and 6.9 x 10^3^ live cells per ml concentration, respectively. All inocula were sprayed onto the abaxial and adaxial side of 5-week old moneymaker tomato leaves following the same procedure as the main passaging experiment. Two experimental controls were included: 6 replicate blank inoculum controls, in which sterile 10 mM MgCl_2_ was applied at the same volume as the treatment plants, and 6 replicate heat-killed controls, in which inocula containing the same concentration of live cells as the treatment plants were autoclaved for 40 minutes to kill the cells. After 3 sequential inoculations and a 1-week incubation period, roughly 1/3 of the leaves (leaf weight range 12.1 – 33.4 g, median = 21.4 g) from the 42 experimental plants were harvested and processed using the same leaf wash protocol as the passaging experiment. The experimental plants were then left to grow for an additional 3 months in the greenhouse, during which time several aspects of host fitness were measured including: number of flowers, number of fruits, fruit weight, number of seeds in the oldest 3 fruits, and the proportion of germinable seeds collected from the oldest 3 fruits.

### Common garden experiment on novel host plants to examine microbiome generalism

To assay microbiome host generalism, we tested the hypothesis that microbiomes with a history of host swapping will more effectively colonize novel hosts and be more resilient to the effects of host filtering. Focusing on the host swapping lines, which all had last colonized tomato hosts, and the conspecifically passaged tomato lines, we inoculated each line onto (i) tomato (*Solanum lycopersicum* var. moneymaker); a host with which the microbiomes share a history of association, as well as onto three novel hosts with no association history: (ii) canola (*Brassica napus* subsp. *napus*), (iii) sorghum (*Sorghum bicolor* var. BTx642), and (iv) corn (*Zea mays* convar. *saccharata* var. *rugosa*); a dicot and two monocots, respectively. The experiment involved a one-time inoculation of plants that were 3.5-4 weeks old. In total there were 6 replicates of each host/microbiome treatment (n = 96 plants), and 6 replicate control plants of each species (n = 24) that were inoculated with sterile 10 mM MgCl_2_. Each entire plant was harvested one week after inoculation and processed in the same way as the passaging experiment. The density of the microbial inocula and the subsequent leaf wash was assessed using both droplet digital PCR (ddPCR) and plate counts of colony forming units (CFUs) of leaf washings.

### DNA extraction, PCR, library preparation, and sequencing

DNA was extracted from every sample of each of the three above-described experiments. One sixth of the total leaf surface microbial community washed from each plant was used for DNA extraction with DNeasy Powersoil Kits (Qiagen). Sample order was randomized to avoid batch effects, and a blank (no sample) control was included in every round of DNA extraction. DNA concentration of each sample was quantified using the Qubit dsDNA HS Assay Kit. Sample DNA (10 μl) was used as template and PCR amplified for 35 cycles at the University of California - Davis Host Microbe Systems Biology Core using the 799F (5’ – AACMGGATTAGATACCCKG – 3’) - 1193R (5’ – ACGTCATCCCCACCTTCC – 3’) primer combination, which targets the V5-V7 region of the 16S rRNA gene, and was designed to minimize chloroplast amplification (35, 36). To further minimize host mitochondrial and chloroplast amplification, peptide nucleic acid (PNA) clamps were added to each reaction (37). Resulting amplicons were diluted 8:1 and were further amplified for 9 cycles to add sample-specific barcodes, then quantified using Qubit, pooled in equal amounts, cleaned with magnetic beads and size selected via electrophoresis on a Pippin Prep gel (Sage Science, USA). The resultant library was then sequenced on the MiSeq (paired-end 300) platform (Illumina, USA).

### Sequence processing

Amplicon sequences were processed using the DADA2 pipeline (38) implemented in the R statistical environment (39), including the packages ShortRead (40), Biostrings (41), and Phyloseq (42). Forward and reverse reads were truncated at 260 and 160 bp, respectively, and quality filtered using the function ‘filterAndTrim’ with default settings (i.e. maxN=0, maxEE=c(2,2), and truncQ=2). Error rates for forward and reverse reads were determined using the ‘learnErrors’ function, and then applied to remove sequencing errors from reads and assign them to amplicon sequence variants (ASVs) using the ‘dada’ function. Paired reads were merged, converted into a sequence table, and then chimeric sequences were removed from the sequence table. Taxonomy was assigned to the remaining ASVs using the ‘assignTaxonomy’ function, which implements the RDP Naïve Bayesian Classifier algorithm with kmer size 8 and 100 bootstrap replicates (43). This taxonomic classification used the Silva (version 138) SSU taxonomic training dataset formatted for DADA2 (44). Chloroplast and mitochondrial sequences were filtered from the ASV table by removing any ASVs with a taxonomic assignment of ‘Chloroplast’ at the Order level or ‘Mitochondria’ at the Family level, respectively. Lastly, we applied the ‘isContaminant’ function (method = prevalence) from the package ‘decontam’ (45) to our samples using our blank (no sample) DNA extractions to identify and remove putative contaminants introduced during DNA extraction.

### Statistical Analysis

All statistical analyses were performed using R version 4.0.3 (39). Community matrices were rarefied to 13500 counts per sample ten times and averaged in order to account for differences in sampling extent across samples. The same rarefying procedure was performed for the specialization and generalism follow up study samples, but to a depth of 31800 and 7900 counts per sample, respectively. Bray Curtis bacterial community dissimilarities were calculated between samples using the ‘vegdist’ function in the vegan package (46). Community structure differences among experimental passage (1-6), host species identity, transmission mode, experimental block, and a host species by transmission mode interaction were assessed using a PERMANOVA on Bray-Curtis distances using the ‘adonis’ function (also in the vegan package), which performs a sequential test of terms and uses the algorithm presented in (47). The order of terms in the model for the field trial was: passage, host, transmission mode, block, and host x transmission, with microbiome line ID specified as strata. The order of terms for the specialization experiment was previous host and experimental block, and for the generalism experiment it was host, previous host, and experimental block. In all cases term order did not impact qualitative conclusions. To assess the change in the relative strength of these factors through time, a PERMANOVA was performed for each of the six passage time points separately. In such instances of hypothesis testing on subsetted data, *p* values were adjusted for multiple comparisons using the Benjamini–Hochberg procedure with the ‘p.adjust’ function (method=‘hochberg’) in the stats package in base R.

In order to assess phylogenetic patterns in the phyllosphere communities, we constructed a phylogenetic tree of all ASVs in the community matrices of the three sequence datasets, which included 8395 ASVs. Sequences were aligned using the ‘AlignSeqs’ function in the DECIPHER package (48) using default settings. Next, pairwise distances between sequences were calculated using the ‘dist.ml’ function in the phangorn package version 2.5.5 (49). These distances were then used to construct a neighbor-joining tree using the ‘NJ’ function in phangorn. Lastly, the neighbor-joining tree was used to create a generalized time-reversible with gamma rate variation (GTR+G+I) maximum likelihood tree using the ‘pml’, ‘update’, and ‘optim.pml’ functions in the phangorn package. We calculated the mean pairwise distance (MPD) of taxa in each sample and compared to the MPD of a community randomization null model (species.pool) to calculate the standardized effect size (SES) using the ‘ses.mpd’ function in the picante package (50). Here a Z-score (the SES of MPD versus the null community) below zero is interpreted as phylogenetic clustering whereby co-occurring taxa are more closely related than taxa drawn at random from the species pool. Z-scores above zero, are taken to indicate phylogenetic overdispersion, where co-occurring taxa are more distantly related than the null expectation. While the use of the 16S rRNA gene for such phylogenetic analyses is reasonable considering it contains both conserved and variable regions, there are nevertheless limitations on our ability to capture functional and/or genomic differences among taxa, especially traits that have been acquired through horizontal gene transfer.

To examine patterns of microbiome network attributes we constructed co-occurrence networks from the community matrices after transforming counts to presence/absence. The function rcorr in the HMisc package (51) was used to generate a Pearson’s correlation matrix with corresponding *p*-values. Following Benjamini-Hochberg correction for multiple comparisons (false discovery rate <0.1), a network was constructed on significant correlations using the igraph package (52), following the procedure described in (53). For univariate data including ASV-level richness, MPD SES, and abundance data, a three-way ANOVA was fit to test for significant effects of the passage, host, transmission mode, and block, with interactions therein. The appropriateness of this procedure was verified by checking for a normal distribution of residuals in the model.

Relative abundance across the complete set of ASVs across plants and transmission groups at the sixth passage for each host plant shows a considerable number of zeros (Supp. Fig. 1, columns 1 & 3). Accordingly, we use a presence/absence model framing to identify taxa that differentially establish under the different transmission treatments. The presence/absence models include a random effect reflecting plant to account for non-independence among observations due to the sampling design. Since we are specifically interested in whether some ASVs were more likely to establish under different transmission modes, we included ASV as a crossed random effect among transmission groups. All models were fitted using the brms package in R (54), with the full model defined as: y ∼ (trans.type|ASV) + (1|plant), using a binomial distribution and logistic regression for presence/absence. The first term is the focal estimate of how transmission mode varies by ASV, and the second term corrects for the fact that different plants may be differently permissive. See Supp. Fig. 2 for details of model convergence and fit.

**Fig. 2:**
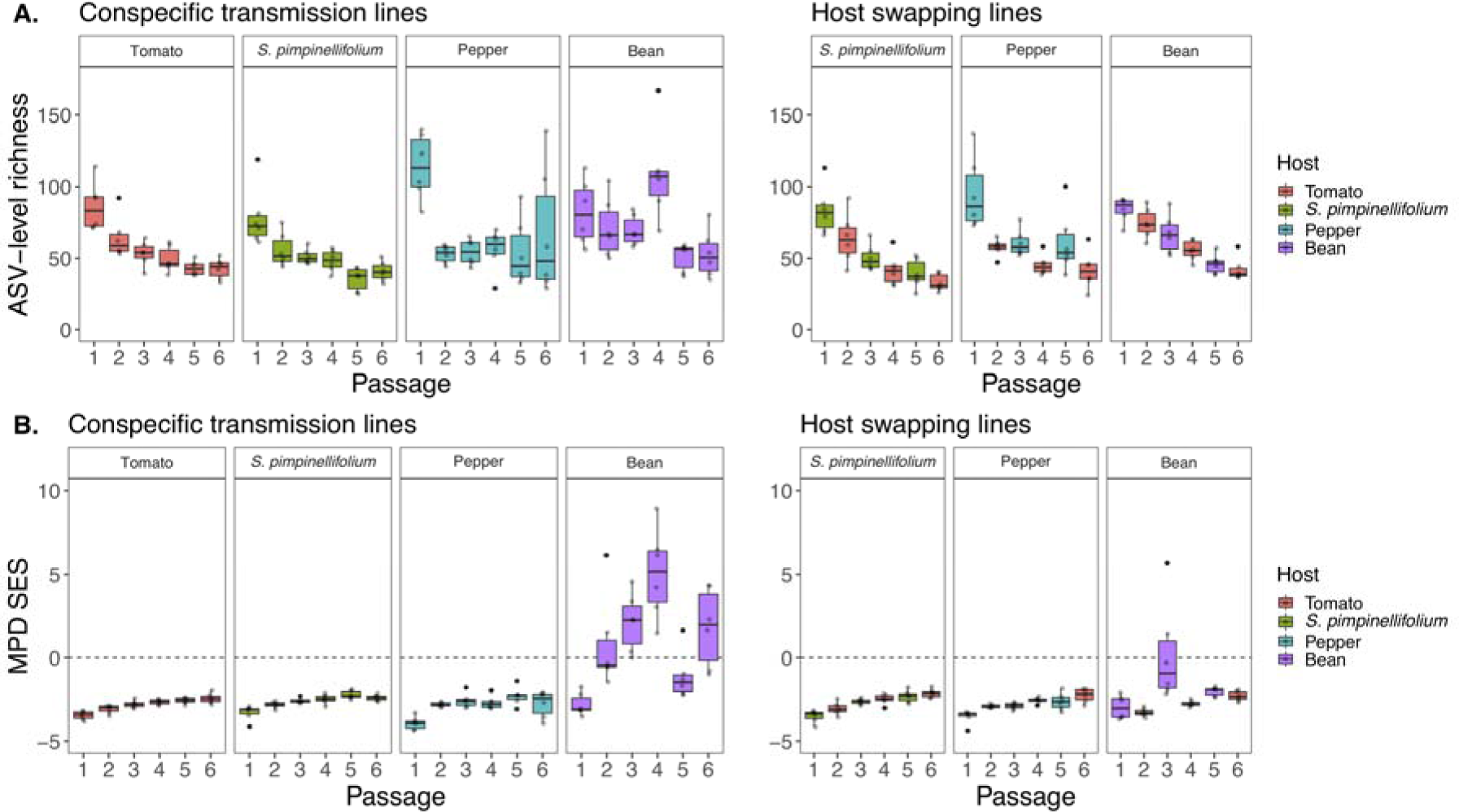
Experimental passaging, host selection, and transmission mode shape phyllosphere microbiome richness and phylogenetic structure. A) ASV-level richness (y-axis) decreases over 6 experimental passages (x-axis). Panels on the left represent the conspecific transmission treatments, panels on the right represent the host swapping treatments, with box colored by host species identity. B) Phylogenetic clustering, represented by the standardized effect size of the mean pairwise distance (MPD SES, y-axis) of conspecific (left panel) and host swapping (right panel) microbiome transmission treatments. SES = (MPD_obs_ - MPD_null_)/SD(MPD_null_). Values below 0 (indicated by dashed line) suggest phylogenetic clustering where co-occurring taxa are more closely related than the null expectation (where taxa are drawn at random from the species pool), while values above 0 suggest phylogenetic overdispersion where co-occurring taxa are more distantly related than null expectation.

## Results

After sample processing and DNA extraction, all samples from the passaging experiment (n = 300) yielded high quality sequences. The dataset contained 7,452,252 observations of 3227 ASVs, 1842 of which had >10 occurrences, and 539 of which had >100 occurrences. The original taxonomic pool from which the inoculum came contained a minimum of 550 ASVs (at detection levels) from Davis, and 139 from Encinitas, with 40 ASVs shared between the sites. Microbiomes from the blank inoculum control plants meant to survey taxa arriving from the greenhouse environment collectively contained 1512 ASVs, 1073 of which were detected only once, and 163 of which were detected in the initial inoculum. Throughout the six passages, taxa from the Encinitas (early growing season) collection on average comprised a higher proportion of read counts in the resulting experimental microbiomes than the taxa from the Davis (late growing season) collection (Supp. Fig. 3), and the taxa shared among the two collection sites tended to comprise roughly one quarter of a microbiome’s sequencing reads (Supp. Fig. 4). At the end of the final passage, phyllosphere microbiome densities ranged from 1.3 x 10^5^ to 3.2 x 10^6^ (median = 9.5 x 10^5^) bacterial cells per g leaf material, and did not significantly differ by host or transmission group (Tukey HSD *p* > 0.1).

**Fig. 3:**
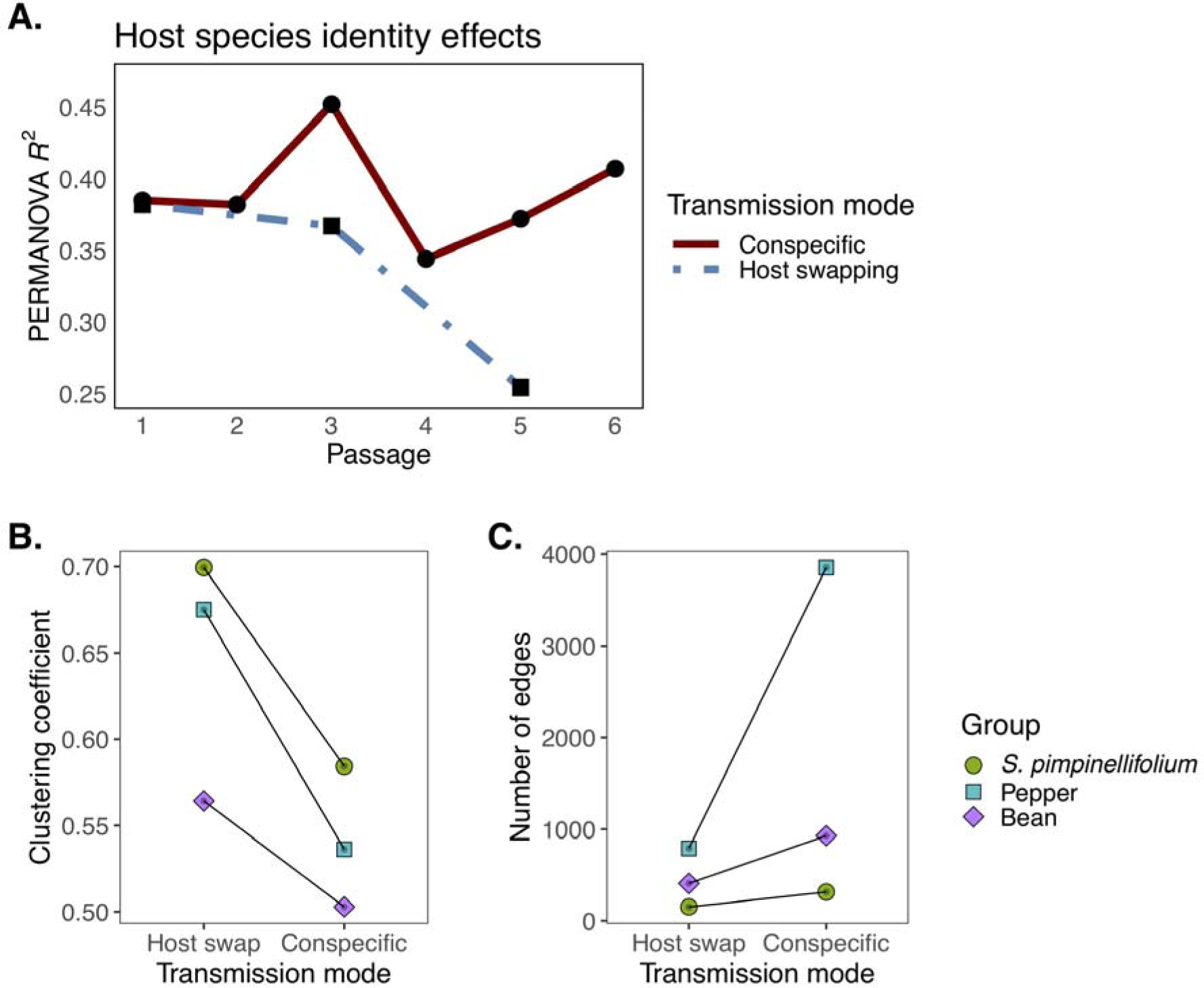
Phyllosphere microbiome composition and co-occurrence network structure are shaped by transmission mode. A) Host species identity effects (y-axis) on phyllosphere bacterial microbiomes are relatively constant over passage time point (x-axis) for conspecific transmission mode (red solid line), while effects decrease in host swapping lines (dot-dash blue line). Y-axis is the PERMANOVA R^2^ value tested on Bray Curtis community dissimilarities with term order: host species identity and block. Statistical significance cutoff is *p* < 0.05, after accounting for multiple hypothesis tests using the Benjamini Hochberg procedure. Note that tomatoes have been removed to make for an even comparison with host swap lines. B) Co-occurrence network clustering coefficient (y-axis) computed from replicate microbiome lines, separated by transmission mode (x-axis) and host group (point shape and color). C) Co-occurrence network number of statistically significant edges (y-axis) computed from replicate microbiome lines, separated by transmission mode (x-axis) and host group (point shape and color).

**Fig. 4:**
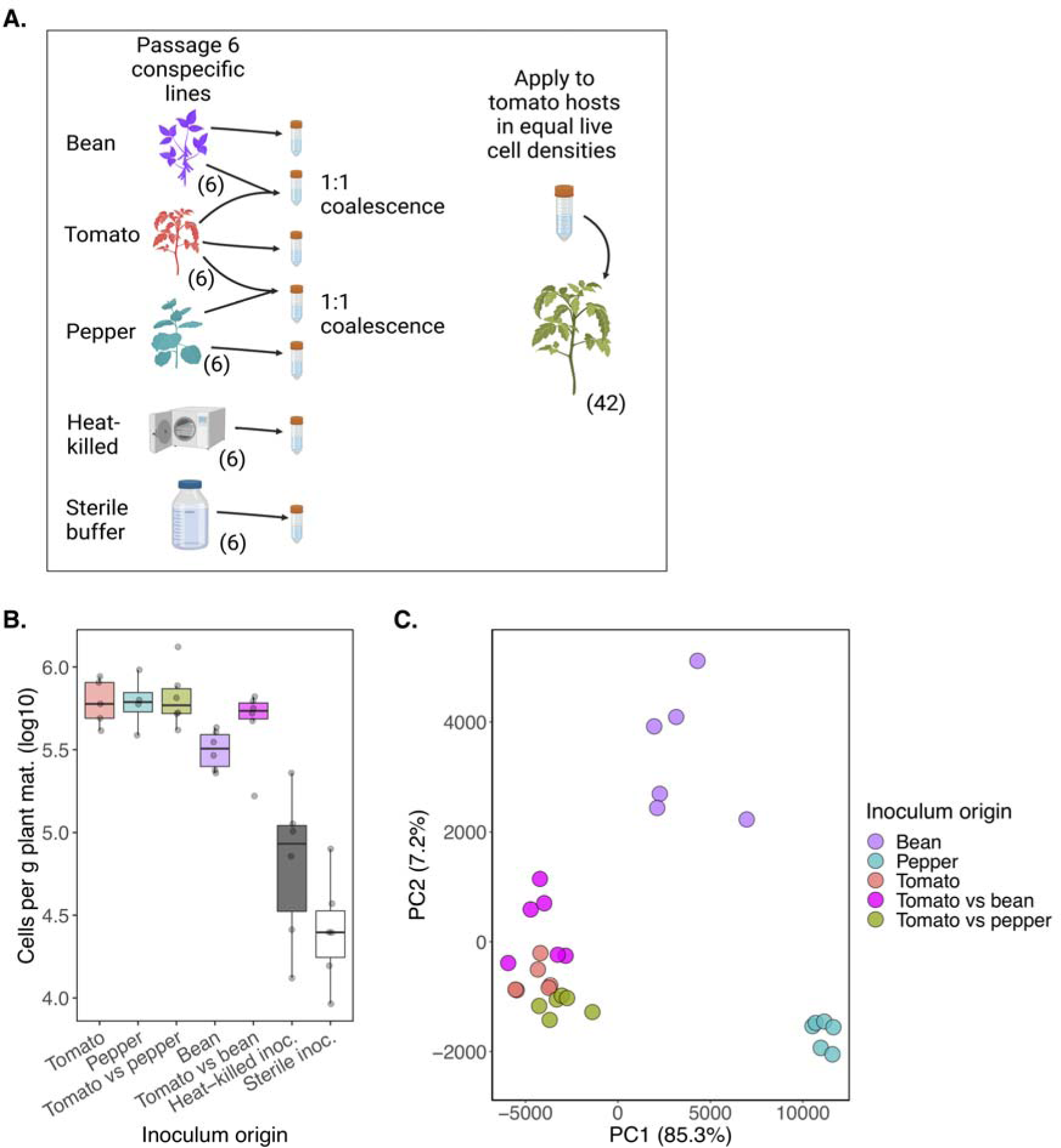
Conspecifically-passaged lines show evidence of host specialization. A) Schematic representation of experimental design in which microbial inocula of different transmission history origins were sprayed on to tomato hosts in equal densities of live cells three times. Inoculum origin is as follows: tomato (conspecifically passaged on tomato), pepper (conspecifically passaged on pepper), bean (conspecifically passaged on bean), tomato vs pepper (community coalescence treatment whereby equal densities of tomato and pepper inocula were combined then sprayed on at the same density as the other treatments), tomato vs bean (community coalescence with tomato and bean inocula), heat-killed inoculum, sterile inoculum (10 mM MgCl_2_). B) Bacterial microbiome densities (bacterial cells per g plant material) on a log10 scale (y-axis), as measured by ddPCR of leaf wash, separated by microbiome treatment (x-axis). C) Principal component analysis of resultant bacterial microbiomes colored by their inoculum origin or coalescence treatment.

### Experimental passage, host species identity, and transmission mode shape phyllosphere bacterial composition

Phyllosphere communities across the 252 treatment samples were dominated by taxa affiliated with the genera *Exiguobacterium*, *Pantoea*, *Sphingomonas*, *Pseudomonas*, and *Klebsiella* (Supp. Figs. 5-7). The PERMANOVA model explaining Bray Curtis dissimilarity among these samples revealed statistically significant effects of passage time point (1-6), the species identity of the host plant, transmission mode (conspecific vs host swapping transmission), experimental block, and an interaction between host species identity and transmission mode (Table 1). In other words, the effects of transmission mode (conspecific or host swapping) depend on the host plant species between which transmission takes place.

**Fig. 5:**
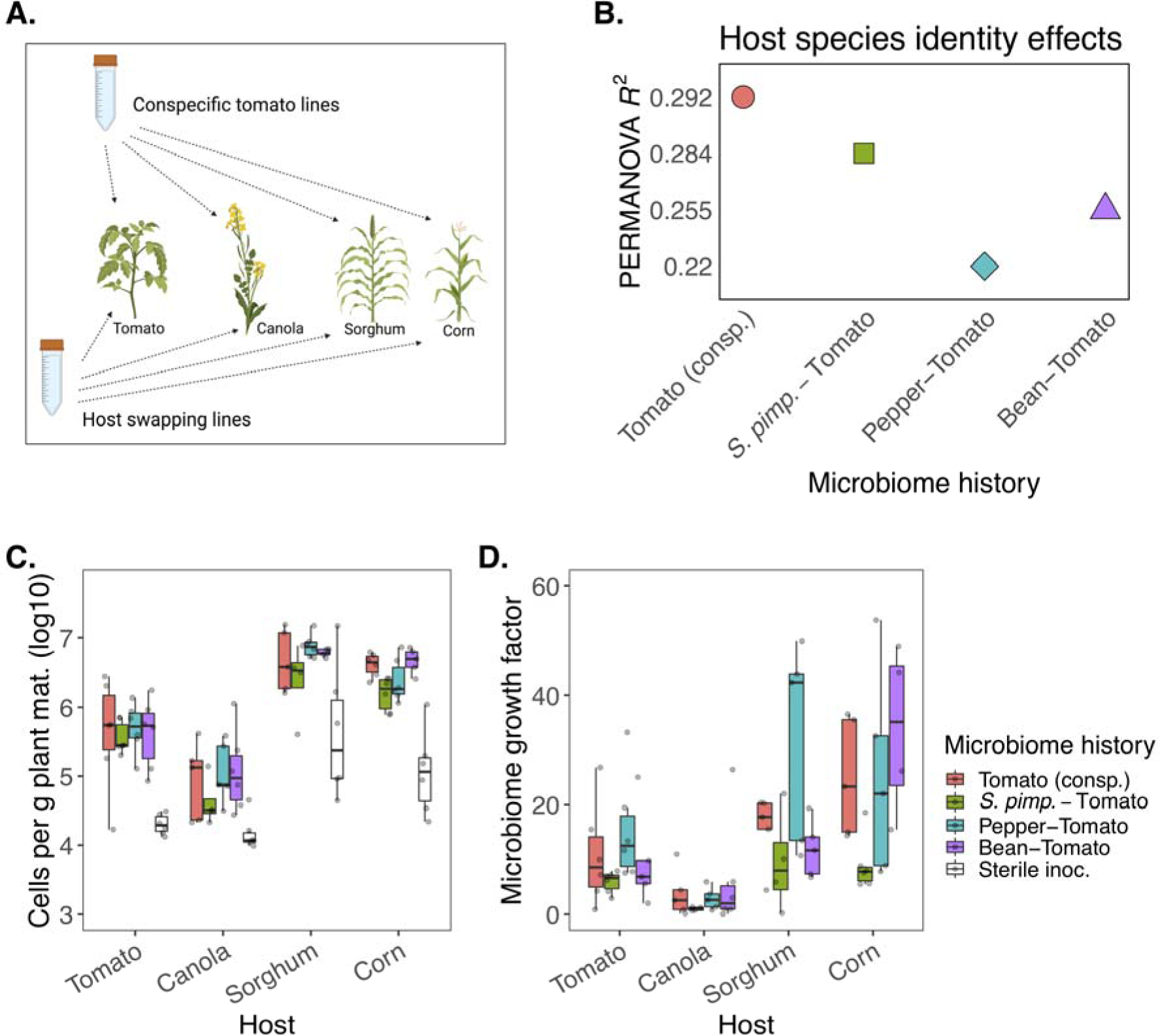
Transplantation of bacterial microbiomes onto novel plant hosts reveals effects of transmission mode, microbiome history, and recipient plant species identity. A) Experimental design schematic illustrating the bacterial microbiome treatment inoculations onto novel host species. Experimental plant species include canola, sorghum, corn, and tomato, and microbiome treatments include conspecifically passaged tomato microbiomes as well as *S. pimpinellifolium* – tomato, pepper-tomato, and bean-tomato host swapping lines. Each treatment/plant combination has 6 replicate plants, and each plant species has 6 replicate sterile inoculum controls. B) Host species identity effects (PERMANOVA R^2^ values, y-axis), representing the distinction among microbiomes by host plant species identity, are lower in the microbiome lines (x-axis) with a history of host swapping. C) Bacterial cells per g leaf material (log_10_ scale, y-axis) is shaped by host species identity (x-axis) and microbiome history (box color). D) Bacterial microbiome growth rate (y-axis) as calculated from the estimated total cells inoculated onto leaf surface divided by the estimated total cells recovered from the leaves one week later. Data are separated by host species identity (x-axis) and colored by microbiome history.

**Table 1:** Phyllosphere bacterial assemblages vary by passage number, host species identity, transmission mode, block, and a species identity-by-transmission mode interaction. Results of a PERMANOVA on Bray Curtis dissimilarities of bacterial communities, with line ID specified as strata. Term order is as follows: passage number, species identity, transmission mode, and experimental block.

| | *df* | *Pseudo-f* | $R^2$ | *p* | *Significance* |
| --- | --- | --- | --- | --- | --- |
| *Passage* | 5 | 36.9 | **0.384** | **0.001** | *** |
| *Host species* | 3 | 11.7 | **0.073** | **0.001** | *** |
| *Transmission mode* | 1 | 1.1 | **0.002** | **0.001** | *** |
| *Block* | 1 | 15.7 | **0.033** | **0.001** | *** |
| *Host species x transmission mode* | 3 | 2.2 | **0.013** | **0.03** | * |

To gain a better understanding of whether the observed effect of passage time point is driven by seasonality, we examined the microbiomes that developed on the seasonality control plants that were treated with the initial inoculum at time points 2-6. A difference in community structure across passage time points was observed in these microbiomes (P2-6, PERMANOVA R^2^= 0.647, *p* < 0.001), however, when compared to the passage 1 tomato microbiomes, there were no significant pairwise differences in composition (pairwise PERMANOVA *p* > 0.05) for any of the passages, suggesting relative similarities in initial host filtering through time from a common pool of microorganisms.

### Microbiome richness steadily declines under host selection

The goal of the passaging experiment was to observe any microbiome community-level responses to host-imposed selection. As such, we predicted continuous loss of taxonomic diversity over time, and expect the differences in taxa lost across independent lines to indicate stochasticity in this process, while the concerted loss across lines to indicate more deterministic processes. Across all lines, ASV richness levels declined over time (F_5, 252_ = 33.56, *p* < 0.001) and were influenced by host species identity (F_3, 252_= 12.49, *p* < 0.001), with a trending but insignificant effect of con-versus hetero-specific transmission (F_1, 252_= 3.01, *p* = 0.08), and no interactions therein (Fig. 2A). Overall, the 6 experimental passages resulted in microbiomes with roughly 2-fold lower richness levels at passage 6 relative to passage 1. At passage 6, all treatments were roughly equivalent in richness, but a trending effect of transmission mode could be detected (F_1, 42_=3.92, *p* = 0.056) wherein conspecifically transmitted groups had slightly higher richness than host swapping groups.

To examine phylogenetic clustering across plant species and transmission modes, we generated a phylogenetic tree comprised of all bacterial ASVs from the experiment to test whether co-occurring taxa within a given microbiome/treatment are more closely related than expected by chance. Evidence for such phylogenetic clustering would consist of a negative standardized effect size of the mean pairwise distance (MPD SES) between pairs of co-occurring taxa. MPD SES values are significantly impacted by passaging time point (F_5, 239_ = 13.24, *p* < 0.001), host species identity (F_3, 239_ = 85.82, *p* < 0.001), transmission mode (F_1, 239_ = 6.41, *p* < 0.05), and a host by transmission mode interaction (F_3, 239_ = 10.17, *p* < 0.001). MPD SES values are consistently negative for the tomato, *S. pimpinellifolium*, and pepper conspecific transmission lines (Fig. 2B, left panel), suggesting that selection for phylogenetically conserved traits in the microbiome had occurred. Lines that were swapped between these hosts were also persistently phylogenetically clustered throughout the experiment (Fig. 2B, right panel). In contrast, lines passaged only on bean exhibited phylogenetic clustering at passage 1, but then trended towards positive values, suggesting phylogenetic overdispersion. The lines that alternated between bean and tomato also trended towards phylogenetic overdispersion when on bean hosts at passage 3.

### Host species identity effects persist through conspecific transmission but decline under heterospecific transmission

To test whether transmission mode impacted the host species identity effects at each passage, we excluded the tomato conspecific lines to enable an even comparison of the 3 host swapping groups and their 3 conspecific transmission counterparts. At each passage time point, conspecific transmission lines exhibited significant host species identity effects, with effect sizes ranging from R^2^ = 0.35 to 0.45, and fluctuating over time (Fig. 3A). In contrast, host species identity effects in the heterospecific transmission lines exhibited a monotonic decrease over the 3 time points when they were on their focal host (i.e., not tomato; P1, 3, and 5). In other words, each time the heterospecific transmission lines transmitted back to their host species of origin, they were found to be less differentiable across host species.

### Previous host association leaves a discernible signal in microbiome structure despite host selection

At passages 2, 4, and 6 when host swapping microbiome lines were all on tomato hosts, we asked whether each line’s previous host association left a discernible signal in the microbiome structure. None of the host swapping lines differed from the conspecifically-transmitted tomato lines at passages 2, 4, or 6 (pairwise PERMANOVA *p_adj._* > 0.1 for all comparisons), suggesting strong effects of host filtering by tomato plants. Despite this evidence for strong host filtering, the effect of previous host species identity was consistently detectable at each of these passages (Passage 2: R^2^ = 0.30, *p_adj._* = 0.003, Passage 4: R^2^ = 0.18, *p_adj._* = 0.04, Passage 6: R^2^ = 0.21, *p_adj._* = 0.04), suggesting that the pool of microbial transplants had been sufficiently shaped by the previous host association to impact the outcome of assembly on tomato hosts.

### Co-occurrence network structure is shaped by transmission mode

We next test the hypothesis that the selection imposed on microbiomes through successive passaging alters the connectivity of microbiome co-occurrence networks, and that the two forms of transmission uniquely shape microbiome networks. We focus specifically on two network attributes: number of edges (i.e. connections between nodes, reflecting how many other ASVs a focal ASV co-occurs with) and clustering coefficient (i.e. how connected the edges of a node are with each other). At passage 6, the resultant microbiome network structures differ by transmission mode. In all three cases the networks of the host swapping transmission groups have a higher clustering coefficient than their conspecific transmission counterparts (Fig. 3B). In other words in these host swapping networks, the ASVs to which a focal ASV (node) is connected tend to be more connected with each other, forming more network clusters. By contrast, in all three cases, the networks of the conspecific transmission groups have a higher number of total edges (i.e. links) than their host swapping counterparts (Fig. 3C). If we divide the edges into positive (persistent co-occurrence) and negative (persistent lack of co-occurrence) associations, we see first that the vast majority of network edges at passage 6 are positive (range of positive edges: 152-3853, median = 404 vs range of negative edges 0-34, median = 0), and secondly that similar to the total edges, the positive edges are consistently higher in the conspecific lines relative to their host swapping counterparts. There are no negative edges for groups involving *S. pimpinellifolium* and pepper, but for groups involving bean, the number of negative edges in host swap lines is 4 compared to 34 in conspecific lines. Thus it appears that the two transmission modes impose different selective pressures on the microbiome co-occurrence networks, with host swapping lines exhibiting more clustering, and conspecific transmission exhibiting more total positive associations.

### More taxa differentially enriched in host swap lines relative to conspecific lines

We next ask whether the two transmission modes enrich for different taxa, for example as a result of host specialization or network restructuring. We use a Bayesian implementation of logistic regression with binomial distribution on ASV presence/absence to assess differential establishment among transmission treatments. At the final time point when we examine the pepper-associated lines, we identify 29 ASVs belonging to 14 bacterial genera (11 families, Supp. Table 1) that are more likely to establish on the host swapping lines, while no ASVs are more likely to establish on the conspecific lines. Similarly for the bean-associated groups, we find 2 ASVs from different families (Supp. Table 1) that are more likely to establish on the host swapping lines and no taxa more likely to establish on conspecific lines. One of these ASVs (belonging to the Erwiniaceae family with no genus-level taxonomy) is enriched in both pepper and bean host swapping groups. This approach does not identify any ASVs that are differentially established in either of the *S. pimpinellifolium* groups. Since the host swapping lines had their final passage on tomato hosts, we further ask whether any differential patterns of presence/absence can be observed relative to the tomato conspecific transmission groups. Indeed, 4 ASVs associated with 3 genera are identified as more likely to establish on tomato conspecific lines relative to pepper host swap lines (Supp. Table 1). Further, 3 of these ASVs are identified as more likely to be on tomato relative to the bean host swap lines (Supp. Table 1). There are no taxa differentially present between *S. pimpinellifolium* host swap lines and tomato conspecific lines and there are no taxa identified as more likely to establish on the host swapping lines relative to conspecific tomato lines.

### Evidence of microbiome host specialization using experimental community coalescence

To test whether conspecific transmission drives host specialization we devised a community coalescence experiment using the conspecifically transmitted tomato, pepper, and bean microbiome lines resulting from passage 6. The six replicate lines for each host species group were combined and used to inoculate a new set of tomato hosts with equal densities of live bacterial cells from each source. Two community coalescence treatment groups were established by mixing equal ratios of live bacterial cells from tomato and pepper groups, or tomato and bean groups (Fig. 4A). Here the expectation is that on a common host, microbiomes will compositionally segregate by their previous host association, and that the coalescence treatments will more closely resemble the tomato microbiomes than the pepper or bean microbiomes due to the competitive advantage the tomato microbiomes would have gained through conspecific transmission.

Bacterial microbiomes of the treatment plants grew to greater densities than that attained on plants treated with the heat-killed inoculum and MgCl_2_ controls (Tukey’s HSD *p* < 0.01), with the exception of plants treated with a bean-derived inoculum, which grew to roughly 5x lower density than other treatment plants and statistically indistinguishable levels as the controls (Tukey’s HSD *p* > 0.1, Fig. 4B). All other treatment plants harbored statistically indistinguishable epiphytic densities (range: 1.7 x 10^5^ – 1.3 x 10^6^ bacterial cells per g plant material, median: 6.0 x 10^5^). The communities on tomato, pepper, and bean lines were significantly distinguishable in composition by their previous host association (Table 2, Fig. 4C), suggesting a lingering differentiation driven by microbiome history. All treatment plants developed microbiomes that were significantly distinguishable in composition from the heat-killed and sterile inoculum controls (Supp. Table 2, Supp. Fig. 8).

**Table 2:**
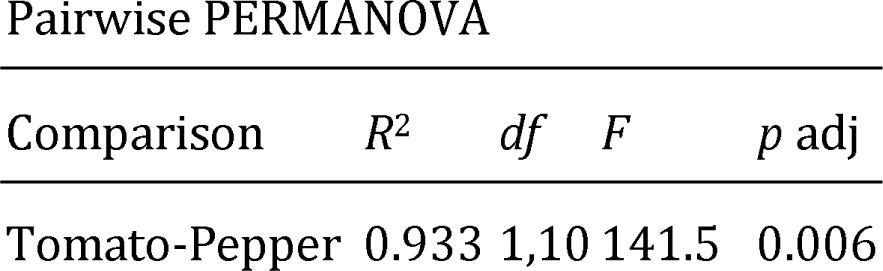
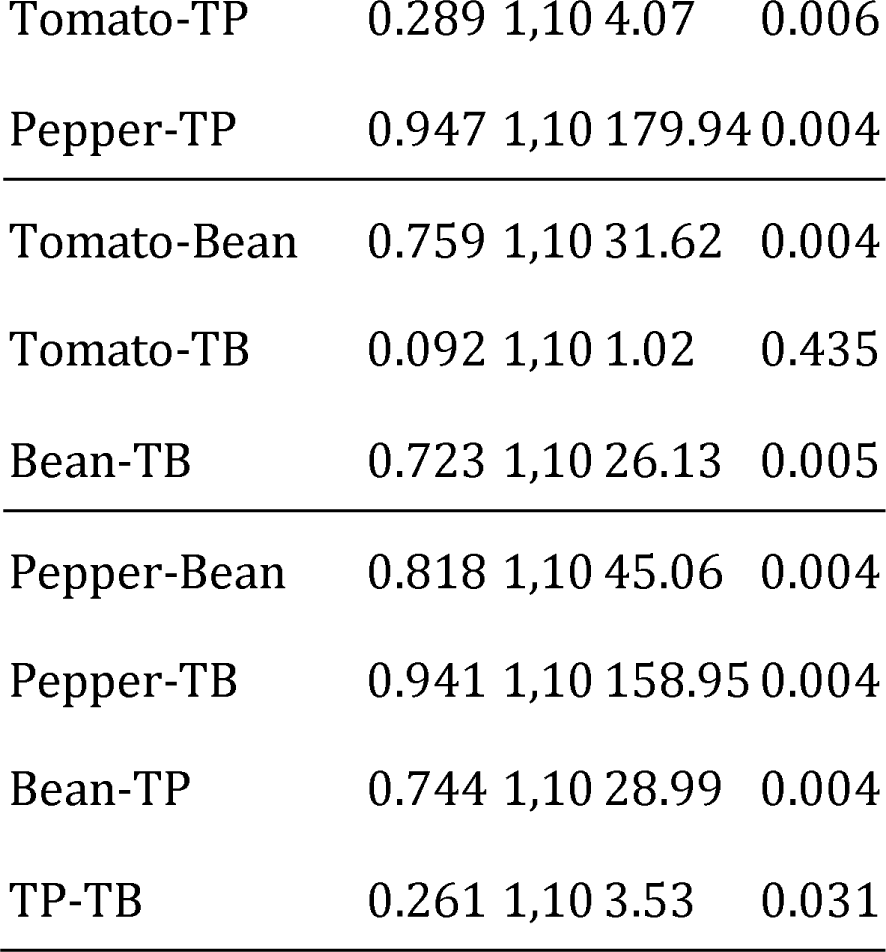
Pairwise PERMANOVA table showing differences of Bray Curtis dissimilarities between microbiomes differing in the host on which they were previously conspecifically passaged for 6 passages, prior to being inoculated on tomato hosts or combined in equal densities as a community coalescence assay. TP = coalescence of tomato and pepper microbiomes, TB = coalescence of tomato and bean microbiomes.

We find support for our hypothesis that tomato microbiome members would come to dominate the pepper and bean microbiomes on a tomato host. The bean-tomato and the pepper-tomato coalescence treatments cluster more closely with the tomato group than the bean or pepper groups, respectively (Fig. 4C). Pairwise PERMANOVA on Bray Curtis dissimilarities reveals that the bean-tomato and tomato microbiomes were statistically indistinguishable (Table 2). The pepper-tomato coalescence and the tomato microbiomes are statistically discernible, but the effect size of these differences is smaller than that between the pepper-tomato and the pepper microbiomes (R^2^ = 0.29 vs R^2^ =0.95, Table 2), suggesting a trend towards dominance of the tomato microbiome.

Since repeated transmission among conspecific hosts has the potential to enrich for host beneficial or deleterious microbial taxa, we monitored the experimental plants for several indicators of health and fitness over a subsequent 3-month period. At weekly intervals over 5 weeks, flower counts increased and then plateaued around week 4, while fruit counts persistently increased (Supp Fig. 9A, B). In both cases during this period, microbiome treatment impacted the total numbers of these organs, driven by bean microbiome-treated plants yielding higher numbers of fruits and flowers than both pepper microbiome-treated plants and heat killed inoculum control plants (Tukey’s HSD *p* < 0.05). At the end of the 3-month period, neither the cumulative production of fruit nor total fruit weight was significantly different among microbiome-treated plants or controls, but the plants treated with the tomato-bean coalescence microbiome had somewhat higher numbers of fruit (Tukey’s HSD *p* = 0.13) and trended towards greater total fruit weight (Tukey’s HSD *p* = 0.07) than plants treated with the heat-killed inoculum (Supp Fig. 10A,B). The average weight of tomatoes did not differ by treatment or from the control (*p* > 0.05, Supp Fig. 10C). Finally, the total number of seeds from the oldest 3 ripe fruits of each plant did not significantly vary among treatments (Supp Fig. 10D), nor did the proportion of seeds that could germinate (Supp Fig. 10E). Thus while experimental microbiomes differ in their ability to establish and flourish on a given plant species, we find little to no evidence for effects on host plant health or fitness.

### Interrogation of the effects of host swapping in novel host environments

We next asked how transmission history would alter the outcome of assembly on a set of novel plant hosts. Each host swapping line was sprayed onto replicate canola, corn, sorghum, and tomato hosts, and then compared to the tomato conspecific transmission lines sprayed onto the same set of host species (Fig. 5A). Compositional variation among resultant microbiomes is explained by both recipient host species identity (PERMANOVA R^2^ = 0.181, *p* < 0.001) and microbiome history (R^2^ = 0.117, *p* < 0.001), with an additional effect of experimental block (R^2^ = 0.080, *p* < 0.01) but no significant interaction between these factors. When we substitute transmission mode (conspecific vs host swapping) for microbiome history, we also see an overall effect (R^2^ = 0.025, *p* < 0.05). Pairwise PERMANOVA reveals that all host comparisons are significantly different in microbiome composition (*p_adj._* < 0.05), and that all microbiome history comparisons are significantly different (*p_adj._* < 0.05), with the exception of tomato conspecific lines and pepper host swap lines, which resulted in compositionally indistinguishable microbiomes (*p* = 0.141). The ASV-level taxonomic richness of the resultant microbiomes is significantly higher in the canola host plants (*p* < 0.001) than the tomato, corn, and sorghum hosts, which are indistinguishable from one another.

When we examine microbiome composition based on the transmission history of the inoculum, we see that all three groups from the host swapping lines resulted in microbiomes with weaker species identity effects than the conspecifically transmitted group (Fig. 5B). In other words, the differences between the resultant canola, corn, sorghum and tomato microbiomes were less pronounced when the inocula came from host swapping lines.

To examine the amount of bacterial microbiome multiplication on the host plants we initially quantified the number of bacterial cells in each inoculum and subsequently measured the number of bacterial cells in the resultant microbiomes one week after their establishment on the plant hosts in the greenhouse. Inocula contained 2.2 x 10^5^ to 4.6 x 10^7^ total bacterial cells, and the subsequent microbiomes grew to 6.6 x 10^5^ to 1.7 x 10^8^ total bacterial cells (1.7 x 10^4^ to 1.6 x 10^7^ bacterial cells per g plant material), an average increase of 19.9-fold. All control plants receiving a sterile inoculum developed significantly lower bacterial abundances than individuals receiving a live inoculum (*p* < 0.001). We find that bacterial abundance per gram of plant material is shaped mostly by host species identity (*F_3,71_* = 80.9, *p* < 0.001), but also microbiome history (*F_3,71_* = 2.79, *p* < 0.05, Fig. 5C). Post hoc analysis reveals that canola plants harbor the least abundant bacterial communities (*p* < 0.001), while corn and sorghum harbor the highest (*p* < 0.001). Similar qualitative conclusions regarding host effects of bacterial abundance were reached through culturing of cells in leaf washings (*F_3,71_* = 24.6, *p* < 0.001), except that no microbiome history effect is detected (*p* =0.871) and instead a host-by-microbiome history interaction is observed (*F_9,67_* = 2.1, *p* < 0.05). The range of microbiome growth factor (i.e. fold growth) is 0.07 – 322.8, and is influenced by host plant species identity (*F_3,69_* = 3.4, *p* < 0.05), with canola hosts harboring the lowest fold increase and corn and sorghum harboring the highest (Fig. 5D).

The taxa from the initial passaging experiment that were identified as enriched in the host swapping lines not only persist in the microbiomes of these novel hosts, but collectively they comprise a majority of the sequencing reads on these plants (79.4 ± 0.15%). When we transform these relative abundances by the ddPCR-inferred absolute abundance estimates of each of the microbiomes, we see that the host swap enriched taxa collectively grew to a range of 6.9 x 10^3^ to 1.5 x 10^7^ (median= 9.0 x 10^5^) cells per g plant material.

Lastly, to examine the resilience, or fidelity, of microbiome transplantation, we calculated the Bray Curtis dissimilarity of the resultant microbiomes on these novel host plants to the samples from which the inocula were derived. This level of divergence was significantly shaped both by the host species identity of the recipient plant (F_3,89_ = 8.21, *p* < 0.001), and the history of the microbiome being inoculated (F_3,89_ = 4.14, *p* < 0.01). When we substitute transmission mode for microbiome history we also see a significant effect on divergence (F_1,91_ = 3.95, *p* < 0.05) in which conspecifically passaged microbiomes diverge more than that of host swapping microbiomes. Post hoc analysis reveals that the communities assembled from conspecifically passaged tomato lines diverged more from their inoculum than the host-swapped *S. pimpinellifolium* lines (Tukey HSD *p* = 0.02), with no other significant comparisons. The difference among recipient hosts is driven by tomato and sorghum plants harboring less divergent microbiomes than corn (Tukey HSD *p* < 0.001 and *p* = 0.03, respectively). We find no evidence for a host-by-microbiome history or a host-by-transmission mode interaction.

## Discussion

The early stages of microbiome assembly are dramatically shaped by dispersal (55), particularly for microbiomes in relatively open habitats such as the phyllosphere. In the case of host-to-host transmission of microbiota, whether taxa arriving early in development come from related or unrelated hosts is likely to be a critical determinant of not just individual microbiome function but perhaps also the (co)evolutionary trajectory of hosts and microbiomes over generations. Recent research has shown that neighboring vegetation can alter the course of phyllosphere microbiome assembly (24), suggesting that the density and composition of the local plant community are likely key factors shaping adaptation of leaf-associated microbial taxa to their hosts. To better understand the consequences of bacterial transmission between members of the same host species (conspecifics) versus between members of different host species (heterospecifics) on microbiome structuring and/or specialization, we implemented a microbiome passaging experiment where transmission mode was manipulated, and then examined the consequences of these transmission modes during assembly on novel hosts and through experimental community coalescence. Our results suggest that the outcome of phyllosphere microbiome assembly depends on microbiome transmission history, and that microbiomes transmitted among conspecific hosts develop a competitive edge during establishment, consistent with host specialization. We further demonstrate that microbiomes swapped among plant species at each passage become less impacted by plant filtering relative to those passaged consistently on the same host plant.

### Use of successive passaging as a means to examine the effects of host filtering

Successive passaging is a promising way to examine how host filtering shapes the assembly, diversity and stability of microbiomes. Host filtering effects were clearly demonstrated throughout our study, indicating that different subsets of microbial taxa survive and reproduce on each of the plant host species (a result previously shown in a field setting; (24)). Consistent with previous work (34), we see a strong impact of passaging on microbiome composition (Table 1) alongside a roughly two-fold reduction of ASV-level richness over the experiment (Fig. 2A). This suggests that selective pressures, as imposed by the plant host and/or within-microbiome interactions, limit the establishment or persistence of many of the taxa from the diverse starting inoculum. Although all microbiome lines tended to converge on roughly similar richness levels, a trending difference could be observed between transmission modes (*p* = 0.08), with those passaged through heterospecific transmission harboring somewhat lower richness than their conspecific transmission counterparts. This may result from the compounding selective filters of both host plants. Evidence for host-imposed selection was also apparent when we examined phylogenetic clustering. Here, we found that the microbiomes on the solanaceous plants - tomato, *S. pimpinellifolium*, and pepper- were significantly phylogenetically clustered, suggesting that the traits under selection – by host filtering, as well as the experimental conditions - are phylogenetically conserved (Fig. 2B). Interestingly, for bean-associated microbiomes we see a trend away from phylogenetic clustering and towards overdispersion. This observation corroborates similar findings of overdispersion and high neutral model goodness-of-fit measures for bean phyllosphere microbiomes compared to tomato and pepper in the field (24). Together, this suggests that beans may be more permissive hosts than tomato or pepper plants. In further support of this concept, host swapped microbiomes involving bean plants trend in a similar phylogenetically overdispersed direction, suggesting a host-by-transmission mode interaction in selective pressures.

Further evidence of strong host filtering can be seen from the effects of microbiome history, i.e. patterns of microbiome differentiation structured by previous host association, observed in all three experiments. In the initial passaging experiment, each time the host swapping lines were transplanted onto tomato hosts, the resultant microbiomes remained statistically discernible in composition by their previous host association, suggesting that previous host filtering can impact subsequent assembly, likely by altering the species pool on which subsequent selection acts. Similarly when microbiomes were transplanted onto novel hosts, microbiome history effects could be observed in the subsequent microbiome composition, diversity, and abundance (Fig. 5C). Finally when the conspecifically transmitted bean, pepper, and tomato microbiomes were transplanted onto tomato hosts we saw a strong effect of microbiome history that resulted in distinguishable compositions and, in the case of microbiomes with a bean-association history, a lower microbiome density (Fig. 4C). Together these observations demonstrate that transmission mode has a pronounced impact on the process of host microbiome filtering, likely by altering the source pool of microbial propagules.

### Microbiomes passaged heterospecifically exhibit homogenization and host generalism

Previous work with phyllosphere microbiomes passaging on different genotypes of tomato found evidence for reduced host identity effects over the course of microbiome selection (Morella et al 2018). In contrast, we find that host identity effects at the species level are retained over time even as microbiome composition shifts in response to selection. Importantly, this was not true of our host swapping (heterospecific transmission) lines, suggesting that this form of transmission might homogenize microbiomes and/or select for more generalist taxa that resist host selective filters (Fig. 3A). This pattern was further exemplified when we transplanted the resultant microbiomes onto a set of novel hosts. Here, the conspecifically-passaged microbiomes were found to exhibit higher host species identity effects than each of the 3 host swapping lines (Fig. 5B), suggesting that the host swap lines are less impacted by the selective filters of the various hosts. Moreover, our differential presence/absence models identified several taxa in the host swapping groups that were more likely to establish on these novel hosts relative to their conspecific transmission counterparts. These taxa collectively grew to high densities on the novel hosts and comprised a majority of the resultant microbiomes, suggesting they may exhibit host generalism and thereby be less impacted by host filtering. Overall, our experimental results suggest that repeated heterospecific transmission has a homogenizing effect on microbiomes and may favor certain microbial taxa with high engraftability on novel hosts.

### Transmission mode shapes microbiome network structure

In addition to differences in the effects of host filtering, another apparent difference in the host swapping versus conspecific transmission lines is in network structure. The co-occurrence networks constructed from the three independent host swapping microbiome lines consistently exhibited a higher clustering coefficient than their conspecific transmission counterparts (Fig. 3B), meaning that edges to a node (i.e. neighbors) tend to be more connected with each other. One implication of a higher clustering coefficient is that the networks could be more resilient to perturbation, such that the loss of any node would be less disruptive to the overall network due to the higher interconnectedness of edges (56). This observation may shed light on the selective pressures the microbiomes face during heterospecific transmission. For instance, if the host swapping treatments face a broader set of host filters by transmitting across two host species, then the development of a more highly clustered network could provide additional stability. Interestingly, we see that the conspecific transmission lines consistently exhibit a higher number of network edges than their host swapping counterparts (Fig. 3C). This may be afforded by the constancy of the host environment that the conspecific transmission lines face. While the use of ecological networks to infer microbial interactions may be limited at relatively large spatial scales such as an entire plant, or through relative abundance measurements (57), the fact that we see these consistent trends across independent biological replicates provides support for the notion that network attributes are in part shaped by transmission mode. Nevertheless additional fine-scale work to interrogate microbe-microbe interactions and microbiome resiliency resulting from transmission manipulation would undoubtedly reveal additional insights into patterns of microbial interactions.

### Conspecific transmission facilitates host specialization: evidence from experimental community coalescence

Community coalescence (i.e. mixing) has been proposed as a framework to examine community-level competition and functionality (29, 58, 59). Here theory predicts that as communities adapt to local conditions through individual member adaptation, cooperation among individual members, or emergence of higher order strategies, they may develop a local competitive edge and hence be more likely to displace competitively inferior communities (60–62). We implemented this framework to examine whether microbiomes could become specialized, i.e. locally adapted, on a given host species through repeated conspecific transmission. We applied microbiomes with a history of tomato, pepper, and bean conspecific transmission to tomato hosts either by themselves or in 1:1 coalescence (competition) treatments and showed that in both coalescence treatments the resultant microbiomes became dominated by the tomato-associated microbiome (Fig. 4C, Table 2). Interestingly the degree to which the coalescence treatments became dominated by tomato microbiomes exhibited a host phylogenetic trend consistent with phylosymbiosis theory (63); with the microbiomes from bean – a much more distantly related host – becoming more dominated than the microbiomes from pepper – a member of the same family as tomato (Solanaceae). Thus, due to the prior repeated assembly history of the tomato-associated microbiomes, these microbiomes exhibited a “home field advantage” on tomato hosts over the pepper- and bean-associated microbiomes, resulting in competitive dominance. These results could be relevant to the conservation of species that are becoming increasingly rare (32), and thus less likely to receive specialized microbial propagules. Our results may also be relevant microbiome engineering efforts, where vigorous establishment and resistance to invasion are desired characteristics (27).

### Concluding remarks: better understanding the role of transmission in shaping microbiome assembly and function

Transmission mode has long been recognized as a key factor shaping the life history strategies of pathogens and other symbionts. Whether such a paradigm applies to complex assemblages of microorganisms residing in or on hosts has remained relatively overlooked. With increasing recognition of the role microbiomes play in plant health, agriculture, and resilience to climate change, it is imperative that we better understand the processes of microbiome assembly and host specialization. While much of the focus of the plant microbiome literature has been on the role the plant host plays in microbiome selection, our results show that by limiting the microbial species pool from which assembly can take place, transmission mode alters the outcome of assembly. We show that conspecific transmission can confer a competitive advantage whereby microbiomes outcompete invading microbiomes with a different host association history, and that host swapping may homogenize microbiomes but confer a broader host range upon them. Such results shed light on how patterns of host–microbiome co-diversification (and phylosymbiosis) may emerge as a result of consistent associations over evolutionary time. Finally, these results also serve as a promising demonstration that by altering transmission mode, microbiome engineering efforts can be directed towards host fidelity and invasion resistance, or broad-range host generalism with taxa refractory to host filtering.

## Acknowledgements

We thank Adam McCurdy and the other farming staff at Coastal Roots Farm in Encinitas, CA and the UC-Davis student farm for providing access to tomato leaves to serve as initial inoculum. We thank R. Porch for assistance in collecting leaves for the initial inoculum. We thank C. Wistrom and the Oxford greenhouse staff at UC-Berkeley for their support throughout the experiment. Figures 1, 4A, and 5A were created with BioRender.com. This study was funded by the US National Science Foundation, award No. 1754494.

