## Supplementary materials for "Transmission mode shapes host specialization of the phyllosphere microbiome"

**Meyer et al. 2023**

**Supp. Figs. 1-10, Supp. Tables 1-2**

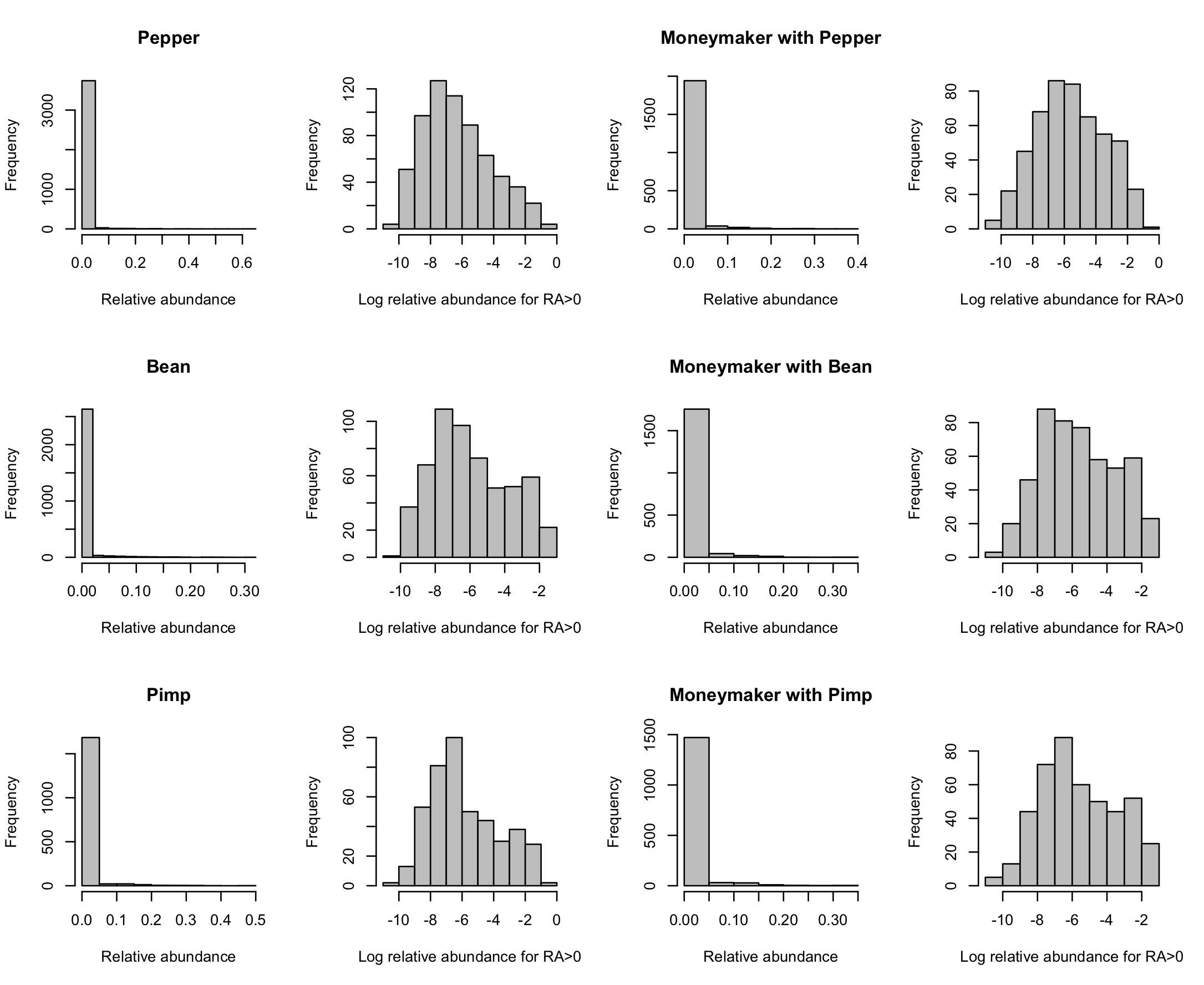

**Supplementary Fig. 1**: Raw distribution of relative abundance (RA) at Passage 6, showing zero-inflation (left hand column, 3152 zeros out of 3804 measurements for pepper; 2191 out of 2760 for bean; and 1311 out of 1752 for *S. pimpinellifolium*); and the distribution of log RA for values greater than 0 (second column); with corresponding observations for tomato moneymaker (1511 zeros out of 2016 measurements for pepper; 1328 out of 1836 for bean; and 1095 out of 1548 for *S. pimpinellifolium*). Plots are separated by their host species and transmission mode (conspecific pepper, host swapping pepper (moneymaker with pepper), conspecific bean, host swapping bean (moneymaker with bean), conspecific *S. pimpinellifolium* (pimp), and host swapping *S. pimpinellifolium* (moneymaker with pimp).

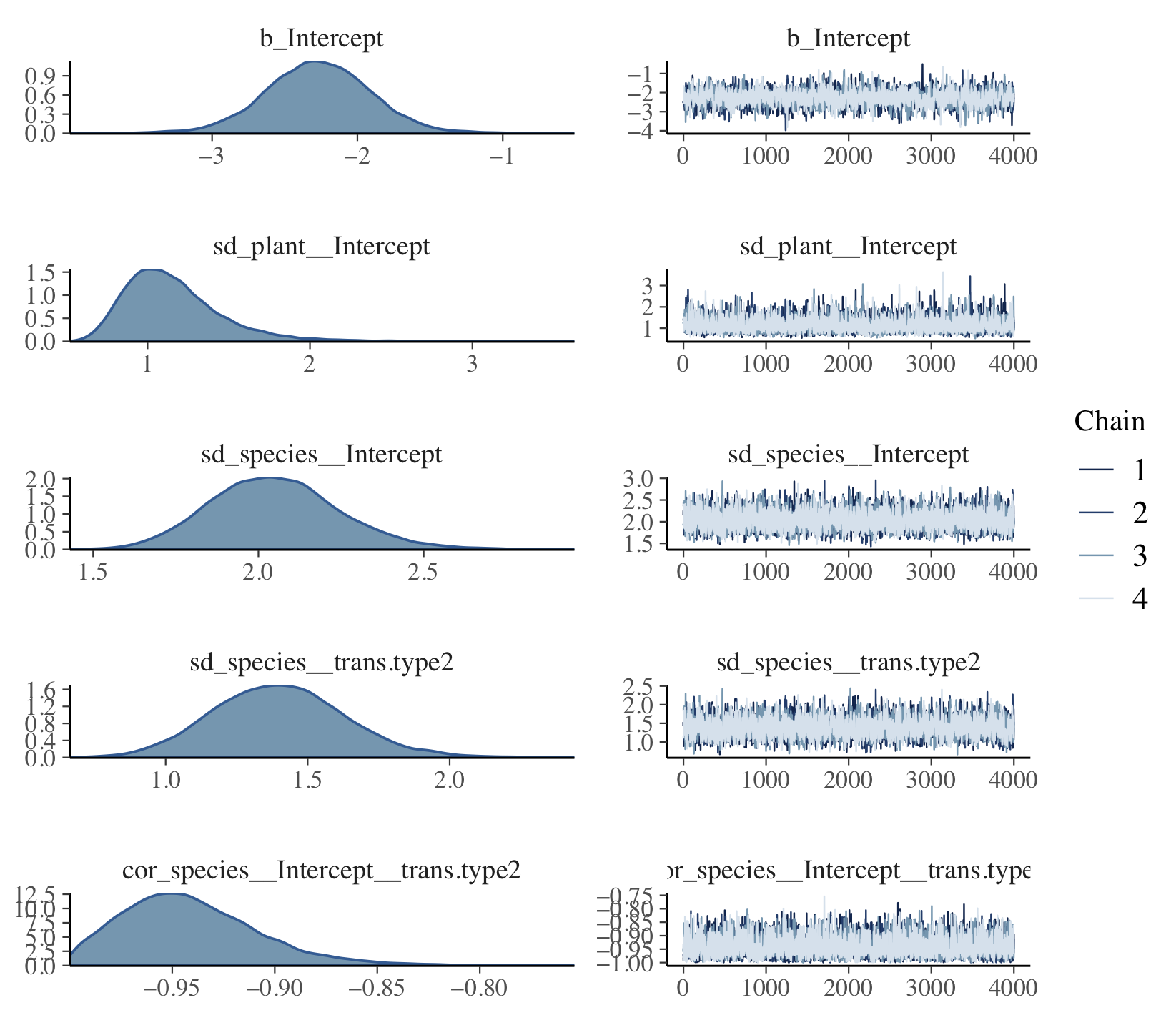

**Supplementary Fig. 2**: Posteriors for the Bernoulli model fitted using the brm package in R, following the model ASV.presence ~ trans.type +(trans.type|species)+(1|trans.type/plant) for pepper plants, others are qualitatively similar. For pepper plants, the Bayesian r^2^ of the model is 0.30; the value of R-hat is very close to 1, indicating that the chains have mixed; for bean plants and *S. pimpinellifolium* plants, the Bayesian r^2^ values of the models are 0.33 and 0.34 respectively; values of R-hat are also very close to 1. For tomato plants with pepper, bean and *S. pimpinellifolium* plants, the Bayesian r^2^ values of the models are 0.39, 0.40, and 0.38, respectively; values of R-hat are also very close to 1.

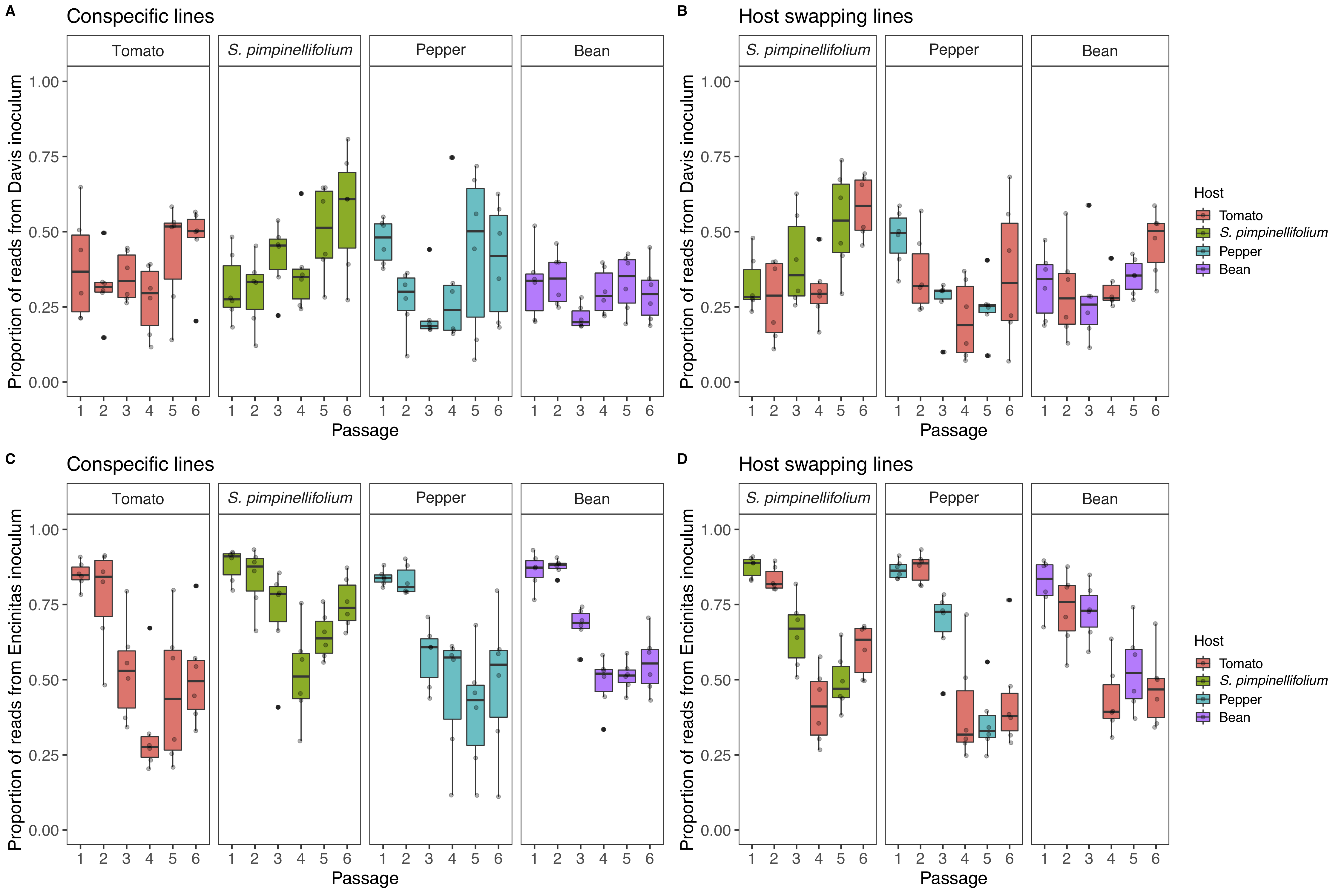

**Supplementary Fig. 3**: The proportion of sequencing reads in each bacterial microbiome that are attributable to taxa detected from the two initial leaf collection sites (y-axis): Davis (A, B) and Encinitas (C, D), California. Microbiome lines in the conspecific transmission treatment are shown in A and C, while heterospecific (host swapping) lines are B and D. For all plots, x-axis is the passage time point.

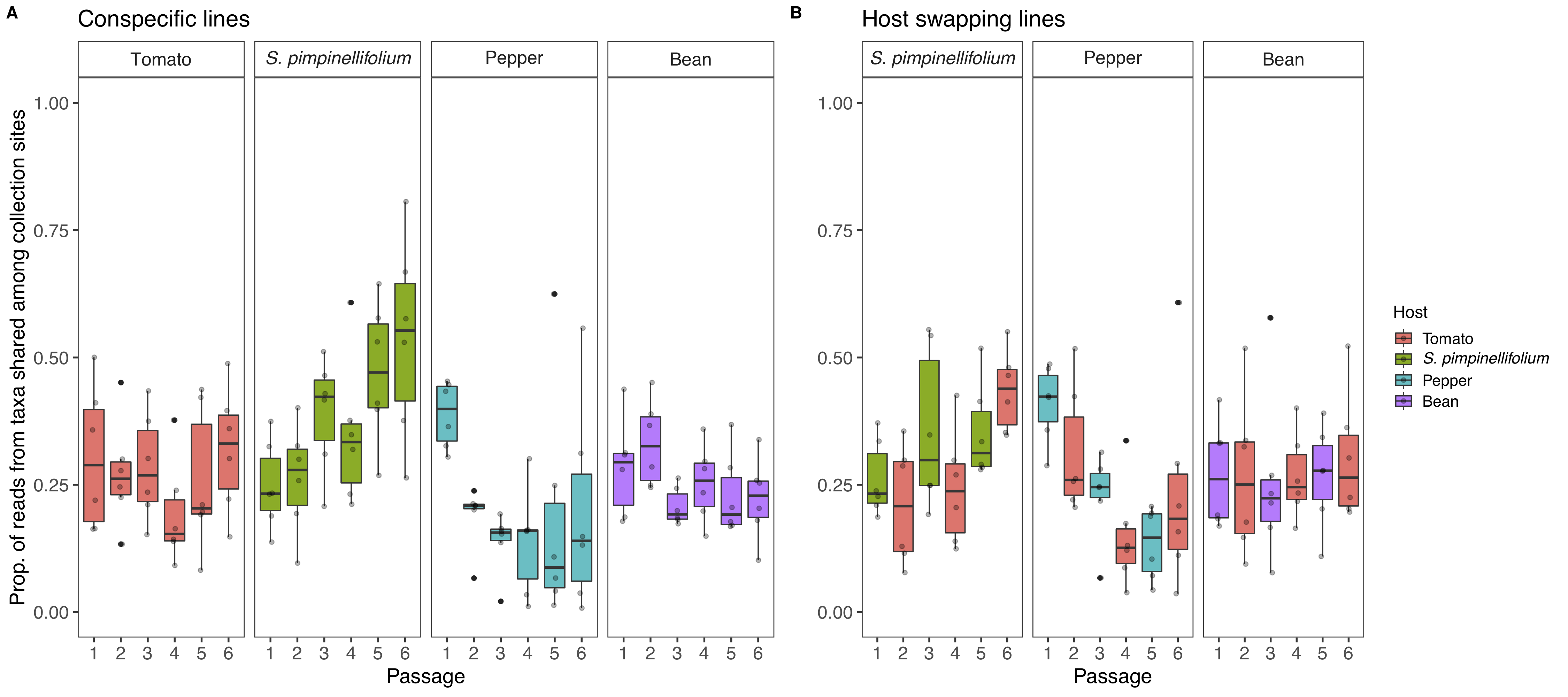

**Supplementary Fig. 4**: The proportion of sequencing reads in each bacterial microbiome that are attributable to the 40 taxa shared between the two initial collection sites (y-axis), faceted by plant host species, with passage time point on the x-axis and panels A and B showing the conspecific and host swapping transmission treatments, respectively.

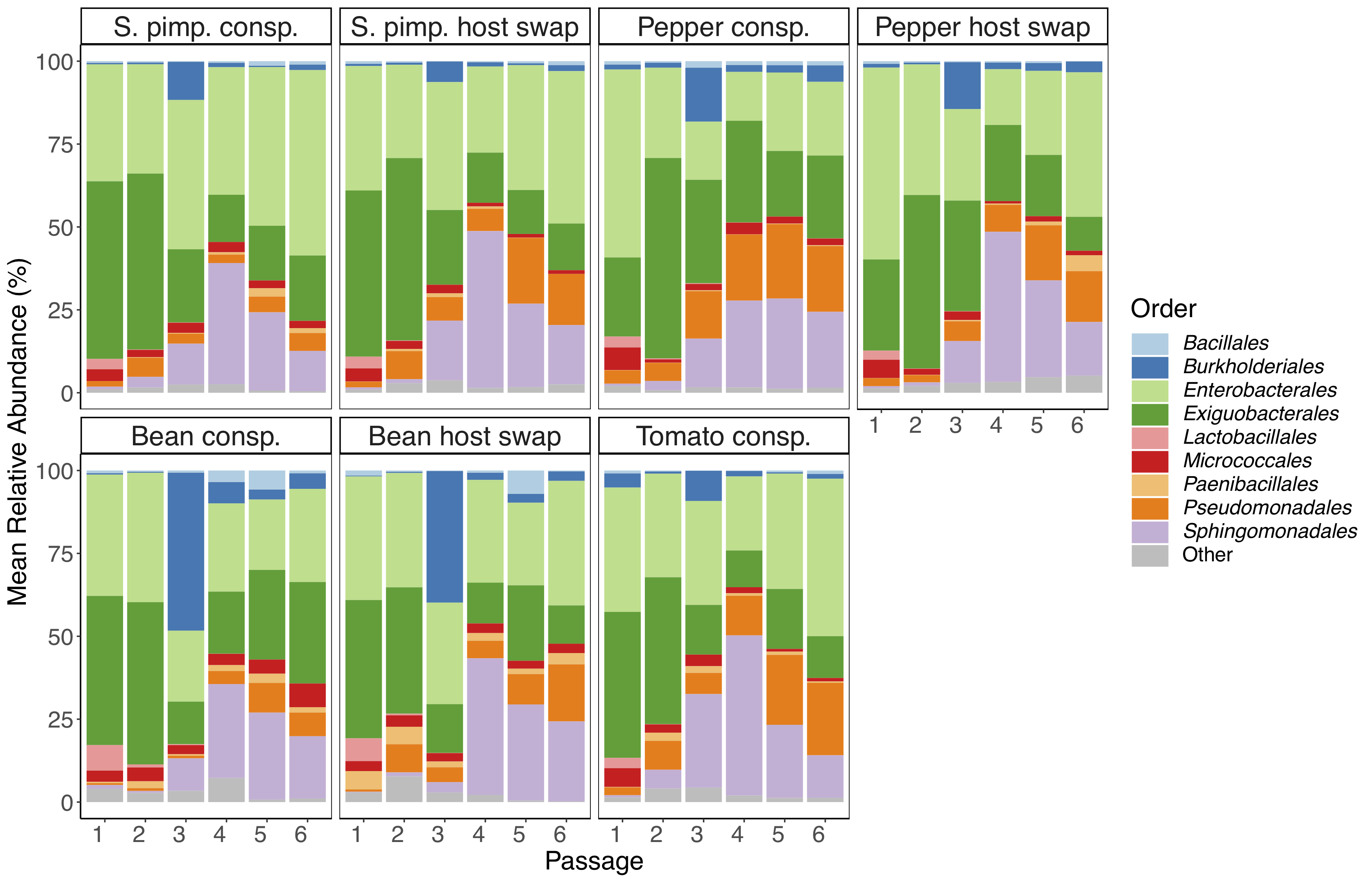

**Supplementary Fig. 5**: Order-level taxonomic bar plots showing relative abundances (y-axis) of epiphytic bacterial microbiomes, faceted by host species (*S. pimpinellifolium*, pepper, bean, and tomato) and transmission mode (conspecific and host swapping) and separated by passage time point (x-axis).

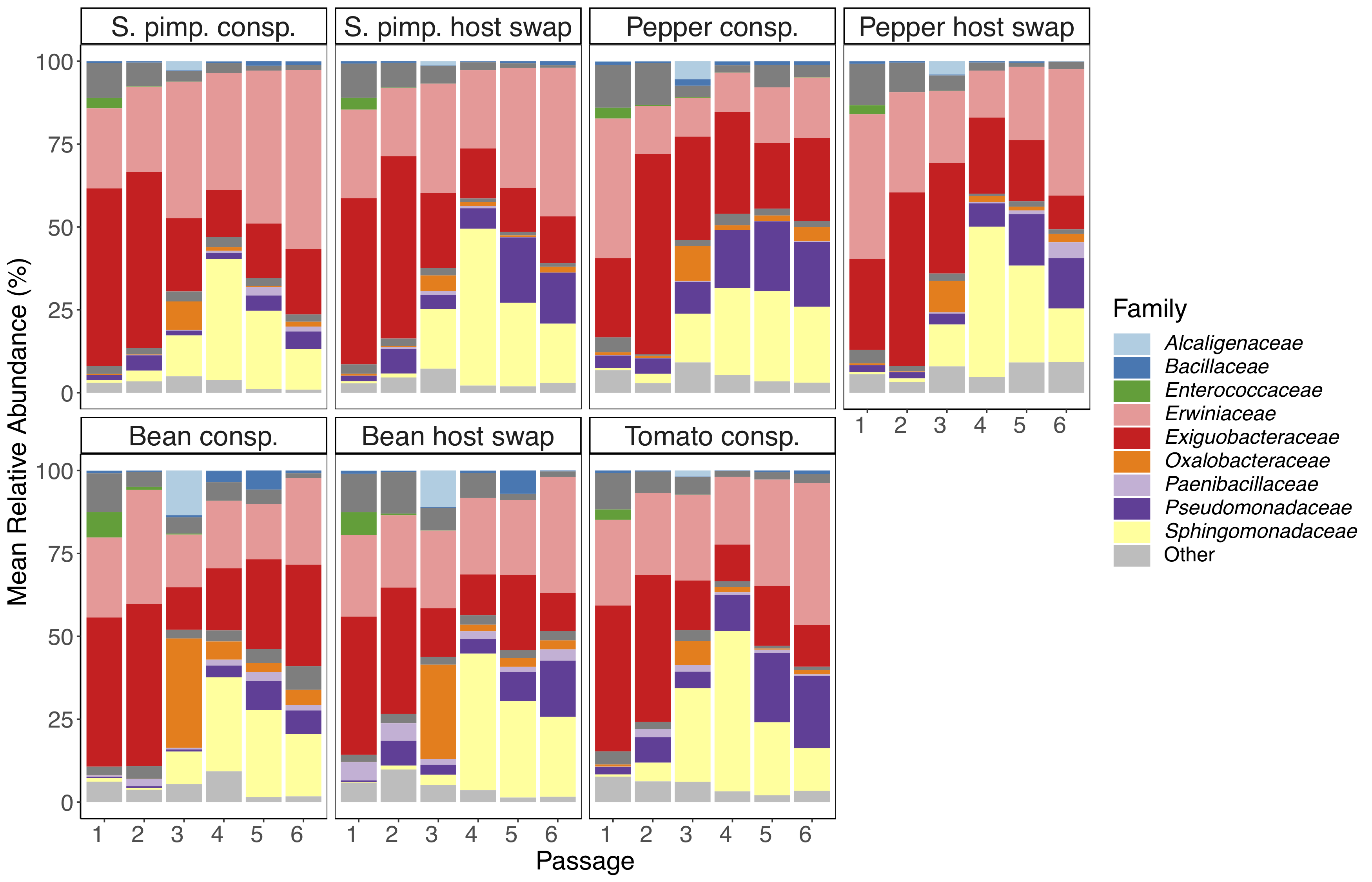

**Supplementary Fig. 6**: Family-level taxonomic bar plots showing relative abundances (y-axis) of epiphytic bacterial microbiomes, faceted by host species (*S. pimpinellifolium*, pepper, bean, and tomato) and transmission mode (conspecific and host swapping) and separated by passage time point (x-axis).

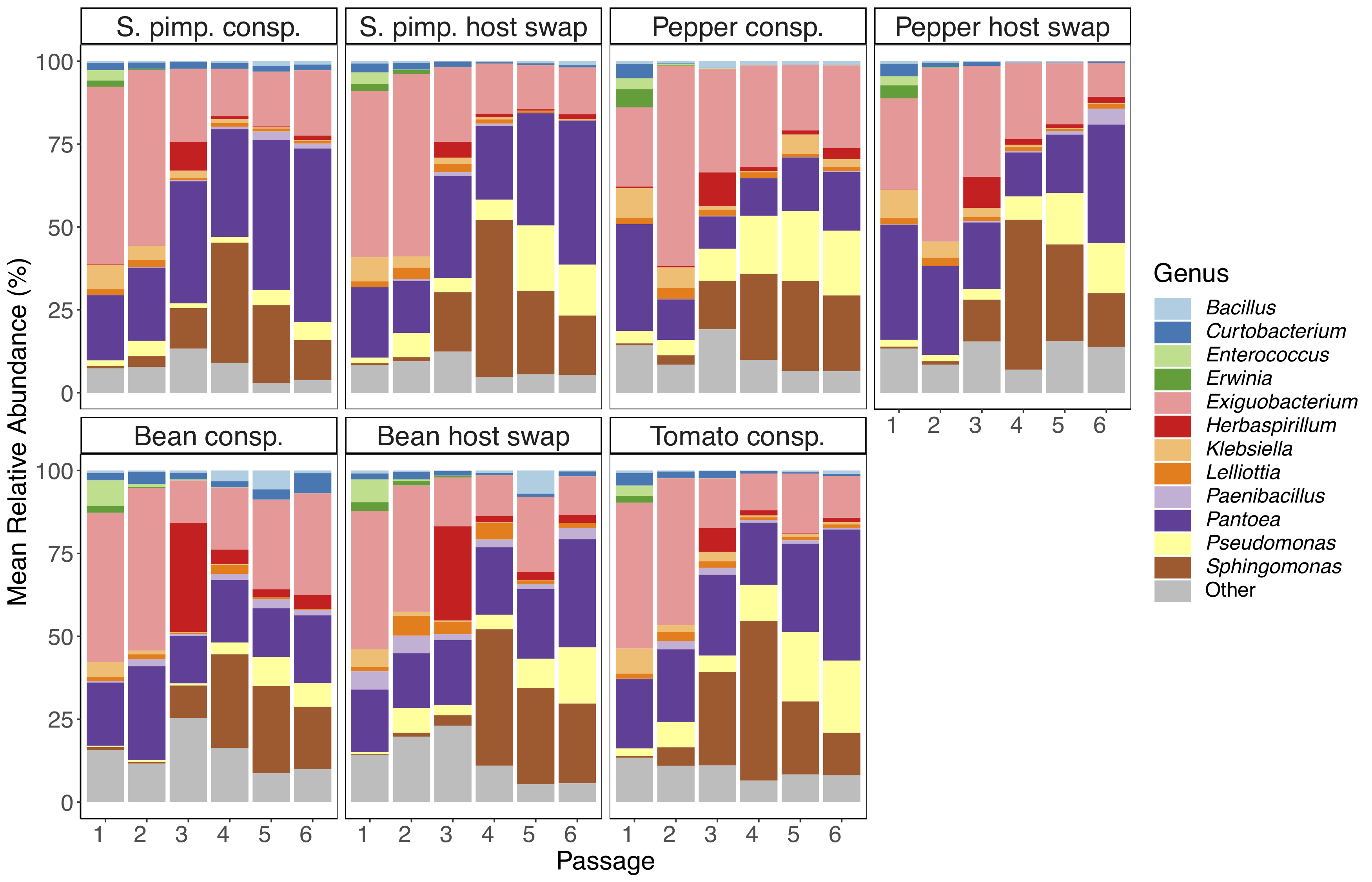

**Supplementary Fig. 7**: Genus-level taxonomic bar plots showing relative abundances (y-axis) of epiphytic bacterial microbiomes, faceted by host species (*S. pimpinellifolium*, pepper, bean, and tomato) and transmission mode (conspecific and host swapping) and separated by passage time point (x-axis).

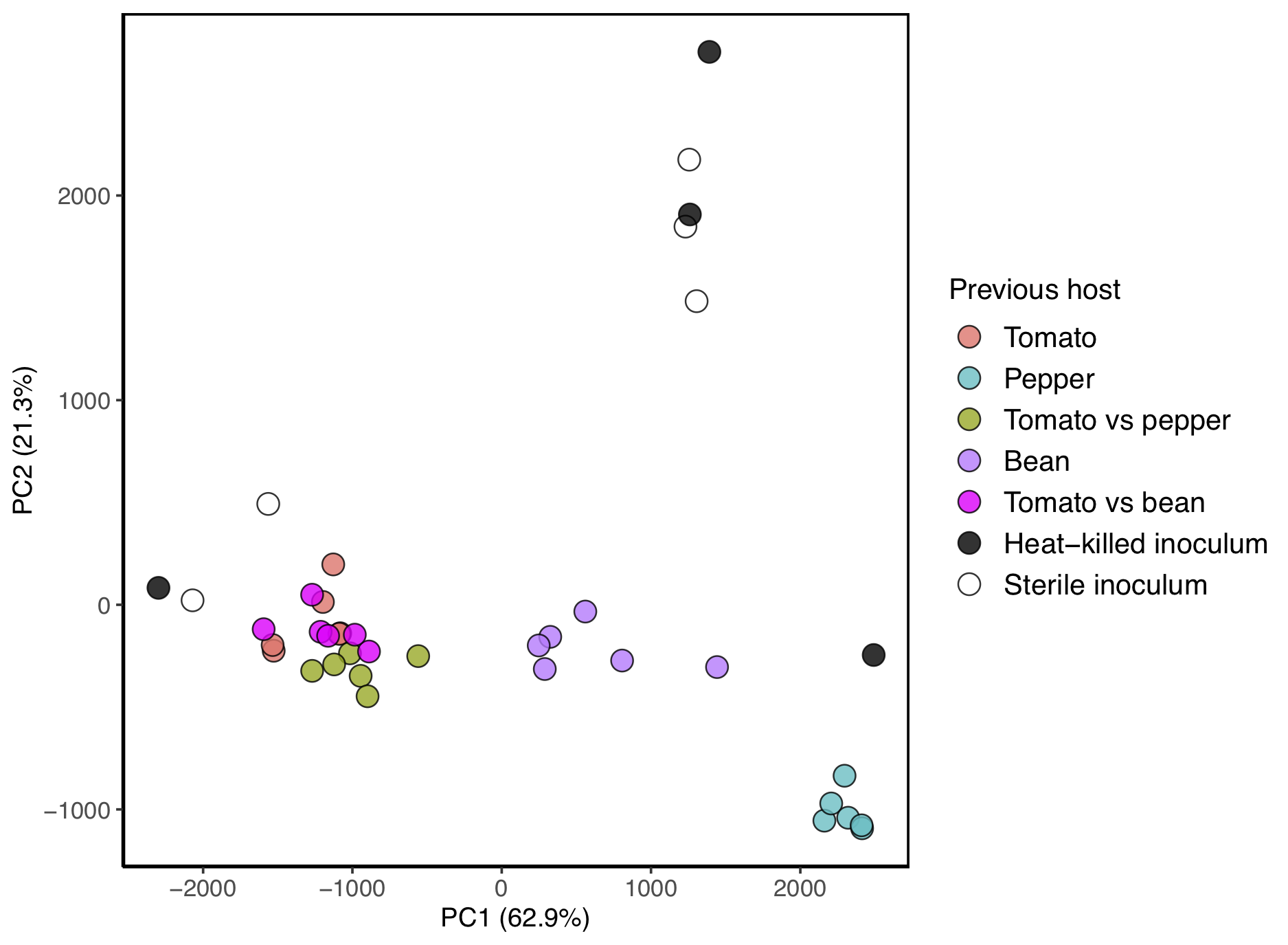

**Supplementary Fig. 8**: Ordination plot showing the position of each experimental bacterial microbiome for the specialization community coalescence experiment along principal components 1 (x-axis) and 2 (y-axis), with percent variance explained in parentheses. Note this is the same ordination as Fig. 4C, but with the heat-killed (black points) and sterile inoculum controls (white-filled points). Points are colored by previous host association, and coalescence treatment, in which two communities were mixed at equal 1:1 ratios.

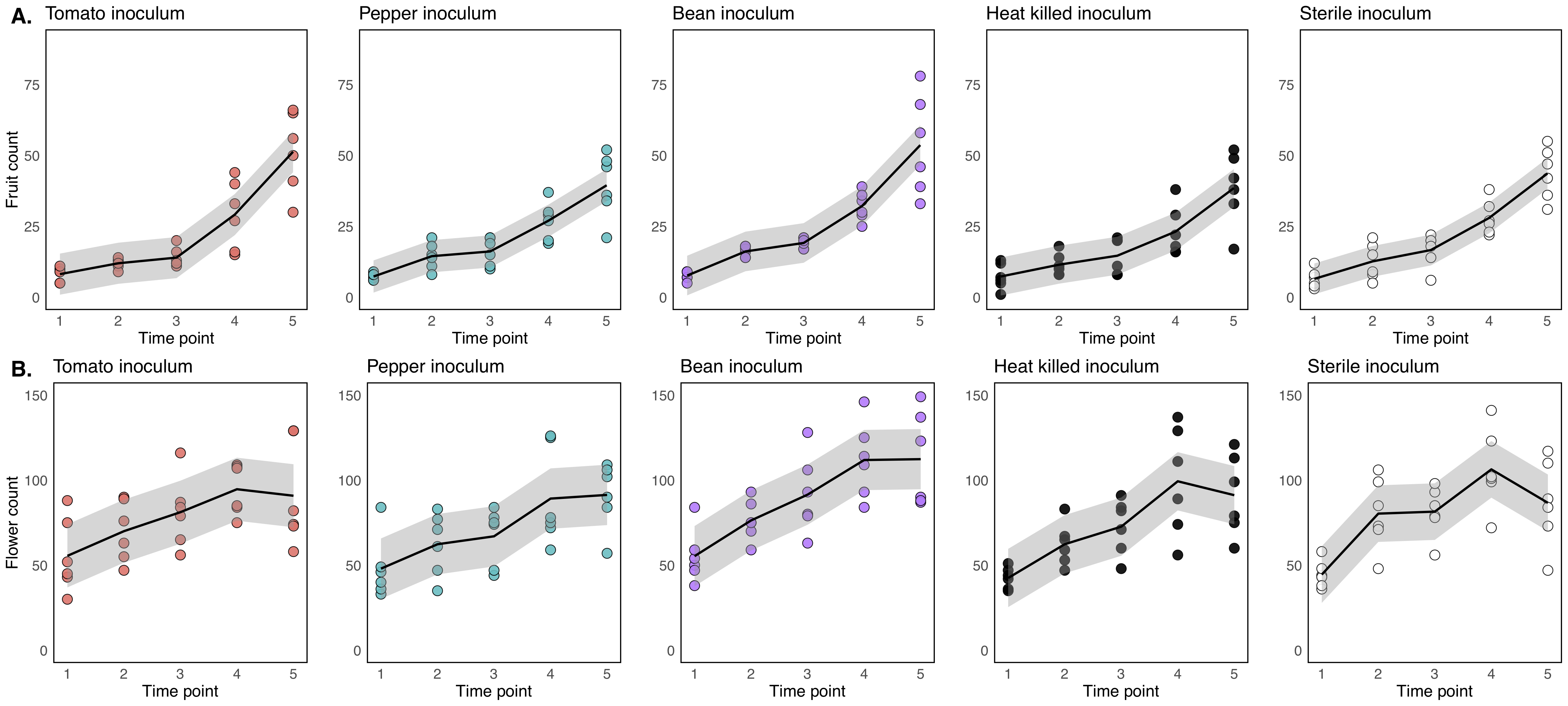

**Supplementary Fig. 9**: Fruit (A) and flower (B) counts (y-axes) collected weekly over a 5-week interval (x-axis) separated by the host or treatment from which inoculum was sourced.

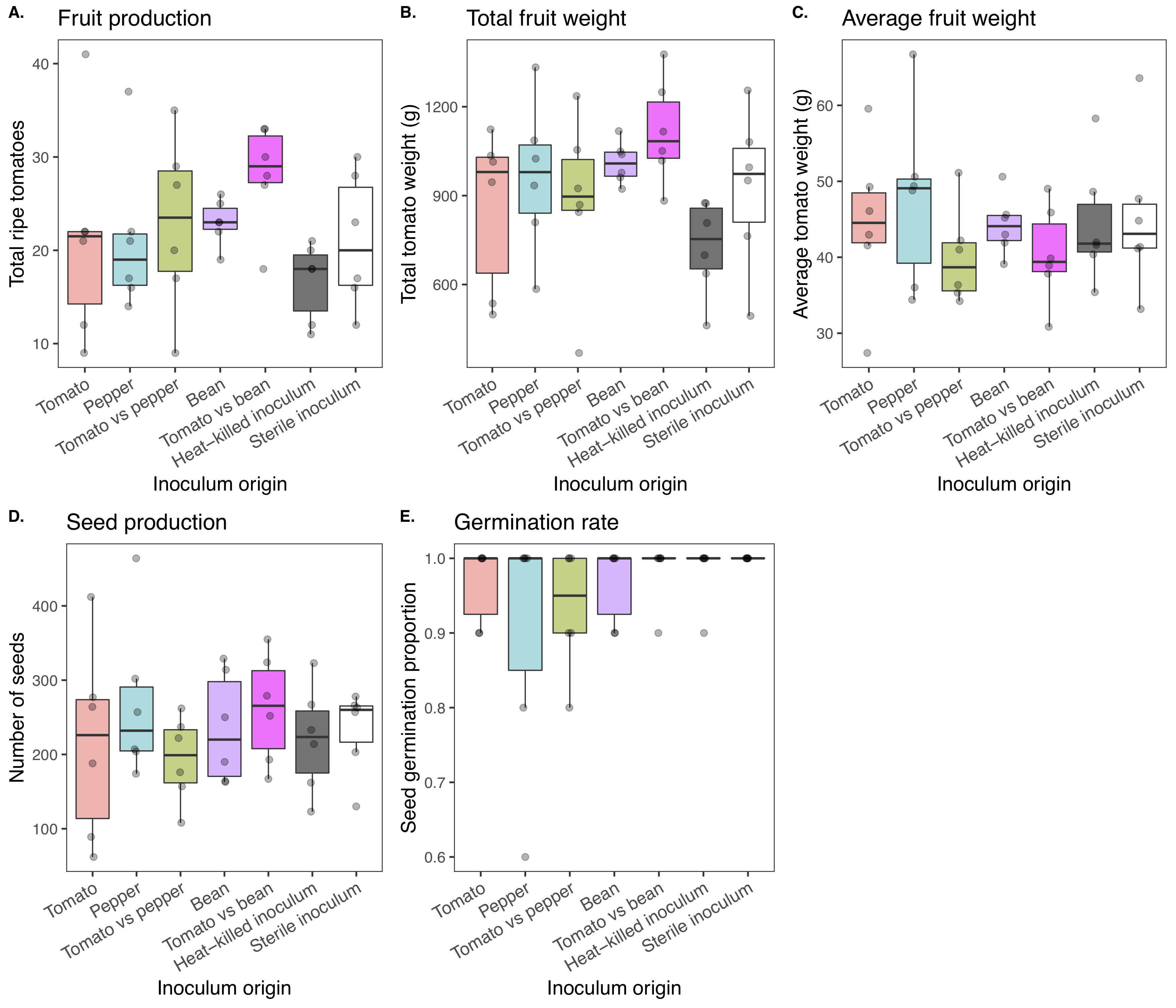

**Supplementary Fig. 10**: Plant health and fitness attributes measured on the microbiome specialization experimental plants over a 3-month period following the survey of microbiome membership. X-axis represents the previous host association of the microbiome. A) Total ripe fruits (y-axis) collected over the 3-month period. B) Total fruit weight (g, y-axis) collected over the 3-month period. C) Average fruit weight (y-axis) of tomatoes collected over the 3-month period. D) The number of seeds from the oldest 3 fruits (y-axis). E) The proportion of germinable seeds (n=10, y-axis) from the 3 oldest fruits.

**Supplementary Table 1**: Taxonomic affiliations of the taxa (ASVs) identified as differentially enriched in a given treatment at passage time point 6.

| **Taxa enriched on pepper host swap lines relative to pepper conspecific lines** | | | | | |  |  |
| --- | --- | --- | --- | --- | --- | --- | --- |
| ASV ID | Kingdom | Phylum | Class | Order | Family | Genus | Species |
| ASV1 | Bacteria | Firmicutes | Bacilli | Exiguobacterales | Exiguobacteraceae | Exiguobacterium | NA |
| ASV2 | Bacteria | Proteobacteria | Gammaproteobacteria | Enterobacterales | Erwiniaceae | Pantoea | NA |
| ASV3 | Bacteria | Proteobacteria | Alphaproteobacteria | Sphingomonadales | Sphingomonadaceae | Sphingomonas | NA |
| ASV4 | Bacteria | Proteobacteria | Gammaproteobacteria | Enterobacterales | Erwiniaceae | Pantoea | NA |
| ASV5 | Bacteria | Proteobacteria | Gammaproteobacteria | Enterobacterales | Erwiniaceae | Pantoea | NA |
| ASV6 | Bacteria | Proteobacteria | Gammaproteobacteria | Pseudomonadales | Pseudomonadaceae | Pseudomonas | NA |
| ASV7 | Bacteria | Firmicutes | Bacilli | Exiguobacterales | Exiguobacteraceae | Exiguobacterium | NA |
| ASV8 | Bacteria | Proteobacteria | Alphaproteobacteria | Sphingomonadales | Sphingomonadaceae | Sphingomonas | NA |
| ASV9 | Bacteria | Proteobacteria | Gammaproteobacteria | Enterobacterales | Erwiniaceae | Pantoea | NA |
| ASV10 | Bacteria | Proteobacteria | Gammaproteobacteria | Burkholderiales | Oxalobacteraceae | Herbaspirillum | NA |
| ASV11 | Bacteria | Proteobacteria | Gammaproteobacteria | Enterobacterales | Erwiniaceae | Pantoea | agglomerans |
| ASV12 | Bacteria | Proteobacteria | Gammaproteobacteria | Enterobacterales | Enterobacteriaceae | Lelliottia | NA |
| ASV15 | Bacteria | Actinobacteriota | Actinobacteria | Micrococcales | Microbacteriaceae | Microbacterium | NA |
| ASV16 | Bacteria | Proteobacteria | Gammaproteobacteria | Enterobacterales | NA | NA | NA |
| ASV17 | Bacteria | Proteobacteria | Gammaproteobacteria | Enterobacterales | Erwiniaceae | NA | NA |
| ASV20 | Bacteria | Proteobacteria | Gammaproteobacteria | Burkholderiales | Burkholderiaceae | Ralstonia | NA |
| ASV22 | Bacteria | Proteobacteria | Gammaproteobacteria | Enterobacterales | Enterobacteriaceae | Klebsiella | NA |
| ASV23 | Bacteria | Proteobacteria | Gammaproteobacteria | Xanthomonadales | Xanthomonadaceae | Stenotrophomonas | NA |
| ASV30 | Bacteria | Proteobacteria | Gammaproteobacteria | Pseudomonadales | Pseudomonadaceae | Pseudomonas | NA |
| ASV37 | Bacteria | Proteobacteria | Gammaproteobacteria | Enterobacterales | Erwiniaceae | NA | NA |
| ASV39 | Bacteria | Proteobacteria | Gammaproteobacteria | Enterobacterales | Erwiniaceae | NA | NA |
| ASV40 | Bacteria | Actinobacteriota | Actinobacteria | Micrococcales | Microbacteriaceae | Curtobacterium | NA |
| ASV52 | Bacteria | Proteobacteria | Alphaproteobacteria | Sphingomonadales | Sphingomonadaceae | Sphingomonas | NA |
| ASV56 | Bacteria | Proteobacteria | Gammaproteobacteria | Pseudomonadales | Pseudomonadaceae | Pseudomonas | NA |
| ASV78 | Bacteria | Proteobacteria | Gammaproteobacteria | Burkholderiales | Oxalobacteraceae | Massilia | NA |
| ASV89 | Bacteria | Proteobacteria | Gammaproteobacteria | Burkholderiales | Oxalobacteraceae | Herbaspirillum | NA |
| ASV93 | Bacteria | Proteobacteria | Alphaproteobacteria | Rhizobiales | Rhizobiaceae | Allorhizobium-Neorhizobium-  Pararhizobium-Rhizobium | NA |
| ASV94 | Bacteria | Proteobacteria | Gammaproteobacteria | Burkholderiales | Oxalobacteraceae | Massilia | NA |
| ASV153 | Bacteria | Proteobacteria | Alphaproteobacteria | Sphingomonadales | Sphingomonadaceae | Sphingomonas | NA |
| **Taxa enriched on bean host swap lines relative to bean conspecific lines** | | | | | |  |  |
| ASV ID | Kingdom | Phylum | Class | Order | Family | Genus | Species |
| ASV18 | Bacteria | Proteobacteria | Gammaproteobacteria | Pseudomonadales | Moraxellaceae | Acinetobacter | NA |
| ASV39 | Bacteria | Proteobacteria | Gammaproteobacteria | Enterobacterales | Erwiniaceae | NA | NA |
| **Taxa enriched on tomato conspecific lines relative to pepper host swap lines** | | | | | |  |  |
| ASV ID | Kingdom | Phylum | Class | Order | Family | Genus | Species |
| ASV19 | Bacteria | Actinobacteriota | Actinobacteria | Micrococcales | Microbacteriaceae | Curtobacterium | NA |
| ASV26 | Bacteria | Proteobacteria | Gammaproteobacteria | Xanthomonadales | Xanthomonadaceae | Xanthomonas | NA |
| ASV44 | Bacteria | Firmicutes | Bacilli | Bacillales | Bacillaceae | Bacillus | NA |
| ASV119 | Bacteria | Actinobacteriota | Actinobacteria | Micrococcales | Microbacteriaceae | Curtobacterium | NA |
| **Taxa enriched on tomato conspecific lines relative to bean host swap lines** | | | | | |  |  |
| ASV ID | Kingdom | Phylum | Class | Order | Family | Genus | Species |
| ASV19 | Bacteria | Actinobacteriota | Actinobacteria | Micrococcales | Microbacteriaceae | Curtobacterium | NA |
| ASV26 | Bacteria | Proteobacteria | Gammaproteobacteria | Xanthomonadales | Xanthomonadaceae | Xanthomonas | NA |
| ASV119 | Bacteria | Actinobacteriota | Actinobacteria | Micrococcales | Microbacteriaceae | Curtobacterium | NA |

**Supplementary Table 2**: Pairwise PERMANOVA table showing differences of Bray Curtis dissimilarities between experimental microbiomes and controls (heat-killed inoculum and sterile MgCl_2_). Experimental microbiomes are labeled by the host on which they were previously conspecifically passaged for 6 passages, prior to being inoculated on tomato hosts or combined in equal densities as a community coalescence assay. TP = coalescence of tomato and pepper microbiomes, TB = coalescence of tomato and bean microbiomes.

| Comparison | *df* | *Pseudo-f* | *R^2^* | *p* | *p_adj_* |
| --- | --- | --- | --- | --- | --- |
| Tomato - Heat-killed | 1,8 | 5.82 | 0.421 | 0.008 | 0.009 |
| Tomato - MgCl_2_ | 1,9 | 5.12 | 0.362 | 0.002 | 0.005 |
| Pepper - Heat-killed | 1,8 | 7.07 | 0.469 | 0.002 | 0.005 |
| Pepper - MgCl_2_ | 1,9 | 10.54 | 0.540 | 0.002 | 0.005 |
| TP - Heat-killed | 1,8 | 5.76 | 0.418 | 0.006 | 0.007 |
| TP - MgCl_2_ | 1,9 | 5.42 | 0.376 | 0.004 | 0.007 |
| Bean - Heat-killed | 1,8 | 5.46 | 0.406 | 0.004 | 0.007 |
| Bean - MgCl_2_ | 1,9 | 6.39 | 0.415 | 0.001 | 0.005 |
| TB - Heat-killed | 1,8 | 5.73 | 0.417 | 0.006 | 0.007 |
| TB - MgCl_2_ | 1,9 | 4.89 | 0.352 | 0.003 | 0.006 |
| Heat-killed - MgCl_2_ | 1,7 | 0.72 | 0.093 | 0.713 | 0.713 |
